# Highly conserved and *cis*-acting lncRNAs produced from paralogous regions in the center of HOXA and HOXB clusters in the endoderm lineage

**DOI:** 10.1101/2020.11.03.366716

**Authors:** Neta Degani, Elena Ainbinder, Igor Ulitsky

## Abstract

Long noncoding RNAs (lncRNAs) have been shown to play important roles in gene regulatory networks acting in early development. There has been rapid turnover of lncRNA loci during vertebrate evolution, with few human lncRNAs conserved beyond mammals. The sequences of these rare deeply conserved lncRNAs are typically not similar to each other. Here, we characterize *HOXA-AS3* and *HOXB-AS3*, lncRNAs produced from the central regions of the HOXA and HOXB clusters. Sequence-similar homologs of both lncRNAs are found in multiple vertebrate species and there is evident sequence similarity between their promoters, suggesting that the production of these lncRNAs predates the duplication of the HOX clusters at the root of the vertebrate lineage. This conservation extends to similar expression patterns of the two lncRNAs, in particular in cells transiently arising during early development or in the adult colon, and their co-regulation by the CDX1/2 transcription factors. Functionally, the RNA products of *HOXA-AS3* and *HOXB-AS3* regulate the expression of their overlapping HOX5–7 genes both in HT-29 cells and during differentiation of human embryonic stem cells. Beyond production of paralogous protein-coding and microRNA genes, the regulatory program in the HOX clusters therefore also relies on paralogous lncRNAs acting in restricted spatial and temporal windows of embryonic development and cell differentiation.

## Introduction

Over the past decade, genome-wide transcriptome analyses revealed a plaetora of noncoding RNAs, that are expressed from a large number of genomic loci. Among those non-coding genes are long noncoding RNAs (lncRNAs), RNA Pol2 products that are longer than 200 nt. Similarly to mRNAs, lncRNAs begin with a 5’ cap and end with a poly(A) tail. To date, thousands of lncRNAs have been reported in different vertebrates (Guttman et al. 2009; Sarropoulos et al. 2019), and it is yet unknown how many of them are functional and what is the full extent of their biological diversity. Many lncRNAs display highly restricted expression profiles during development, potentially allowing them to control gene expression in specific cellular contexts (Mercer et al. 2008; Sarropoulos et al. 2019). Some lncRNAs have been shown to indeed contribute to proper embryonic development (Perry and Ulitsky 2016).

Mouse and human Hox genes are organized in four genomic clusters (HOXA to HOXD) that exhibit a unique mode of transcriptional regulation – temporal and spatial collinearity – the position of the genes along the chromosome roughly corresponds to the time and place of their expression during development. The sequential activation of Hox genes in the primitive streak helps determine the subsequent pattern of expression along the anterior–posterior axis of the embryo (Deschamps and van Nes 2005; Izpisúa-Belmonte et al. 1991). Despite the crucial importance of Hox genes during development (Kmita et al. 2005), the molecular pathways that dictate their collinear expression remain mostly unknown.

Noncoding RNAs are likely to play important roles in Hox gene regulation. For example, Hox clusters encode two conserved miRNAs, miR-10 and miR-196 (iab-4 in *D. melanogaster*), that target some of the Hox genes and help establish specific regulatory programs in the embryo (Hornstein et al. 2005; Mansfield et al. 2004; Tyler et al. 2008). One of the first lncRNAs that has been studied in detail, *HOTAIR*, is produced from the HOXC cluster and was reported to regulate expression of HOXD genes (Rinn et al. 2007). Since this seminal discovery, numerous lncRNAs have been implicated as important in the Hox gene regulation (Casaca et al. 2018). For example, *HOTTIP*, a lncRNA is located at the 5’ end of the HOXA cluster, was shown to control activation of 5’ HOXA genes in *cis* via cooperation with an MLL histone methyltransferase complex and chromosomal looping that brings it into close proximity with 5’ HOXA gene loci (Wang et al. 2011).

The protein-coding genes in the four vertebrate Hox clusters belong to 13 groups of orthologs that can be traced to ancestral clusters that existed before the two rounds of genome-duplication (Hoegg and Meyer 2005). The two conserved microRNA families encoded in the Hox clusters, miR-10 and miR-196, are represented in multiple clusters (Yekta et al. 2008). lncRNAs have been described in each of the four clusters but so far there were no known cases of clear similarity between lncRNAs across clusters. Here, we focus on a pair of lncRNAs that appear to be some of the most conserved lncRNAs produced from the vertebrate Hox clusters – *HOXA-AS3* and *HOXB-AS3*. We provide evidence that it is likely that the production of these lncRNAs precedes the duplication of the ancestral Hox cluster into HOXA and HOXB. Both lncRNAs are expressed predominantly in the embryo, with expression patterns more similar to each other than to nearby protein-coding genes. In the adult, *HOXA-AS3* expression is mostly restricted to tissues of endodermal lineage, and specifically to immature goblet cells and tuft cells. The similar expression of *HOXA-AS3* and *HOXB-AS3* is likely driven by conserved and shared binding sites for CDX transcription factors in the *HOXA-AS3* and *HOXB-AS3* promoters. Using human cell lines and human embryonic stem cells, we show that perturbation of *HOXA-AS3* and *HOXB-AS3* expression results in corresponding changes in expression of HOX-6 and HOX-7 genes. These results suggest co-ordinated and ancient lncRNAs production from central regions of the Hox clusters that plays important *cis*-acting gene regulatory roles in cells of the endodermal lineage.

## Results

### A pair of conserved lncRNAs in the middle of HOXA and HOXB clusters

In human, *HOXA-AS3* transcription starts ~700 nt downstream of the annotated 3’ end of *HOXA5* and it is transcribed antisense to *HOXA5* and *HOXA6*, terminating in the single intron of *HOXA7* (**Fig. 1A**). The region in the mouse genome that aligns to the *HOXA-AS3* promoter is the promoter of *Hoxaas3* (*2700086A05Rik*), which terminates in the intergenic region between *Hoxa6* and *Hoxa7* (**Fig. 1A**). The promoter of *HOXA-AS3* is highly conserved in other vertebrates, but transcripts originating from it are not consistently annotated, likely due to its very restricted expression in adult tissues, as it is expressed predominantly in the embryo (see below). Using available RNA-seq data we could identify homologs for *HOXA-AS3* in opossum and *X. tropicalis* (**Fig. 1A** and **S1**). Transcription of these orthologs, similarly to that of the human *HOXA-AS3*, started ~500 nt downstream of the 3’ end of *HOXA5* and ended in the intron of *HOXA7*.

**Figure 1.**
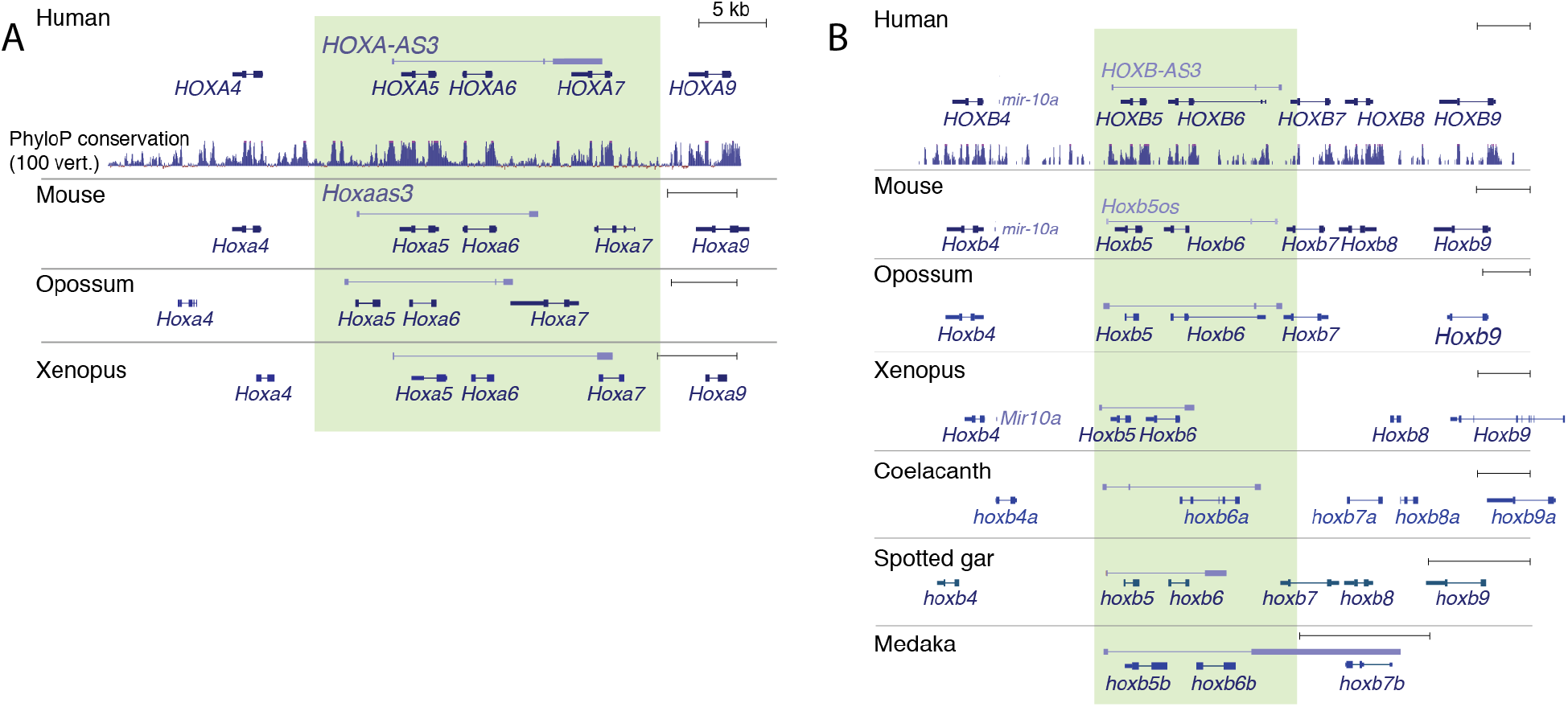
Homologs of *HOXA-AS3* and *HOXB-AS3* in different vertebrate species. Transcript models annotated by Ensembl, Refseq, or PLAR (Hezroni et al. 2015), or manually reconstructed based on RNA-seq data (see **Fig. S1**) for *HOXA-AS3* (**A**) and HOXB-AS3 (**B**) are shown alongside the annotated protein-coding genes in the locus. The lncRNAs are transcribed from the ‘+’ strand and all other genes are transcribed from the ‘-’ strand. The regions of *HOXA-AS3* and *HOXB-AS3* are shaded.

*HOXB-AS3* transcription in human starts ~900 nt downstream of the 3’ end of *HOXB5* and terminates in the intergenic region between *HOXB6* and *HOXB7* (**Fig. 1B**). Presumably because of its broader expression compared to *HOXA-AS3*, homologs of *HOXB-AS3* were readily identifiable in more species. In mouse, it is annotated as *Hoxb5os* (*0610040B09Rik*), and we could identify homologs in opossum, *X. tropicalis*, coelacanth, spotted gar, medaka, and elephant shark (**Fig. 1B** and **S1**). Both *HOXA-AS3* and *HOXB-AS3* show negative PhyloCSF (Lin et al. 2011) scores throughout the locus (**Fig. S2**), and so it is unlikely that they encode highly conserved proteins. Notably, a primate-specific protein has been recently found to be encoded by *HOXB-AS3 (Huang et al. 2017)* (see Discussion).

**Figure S1.**
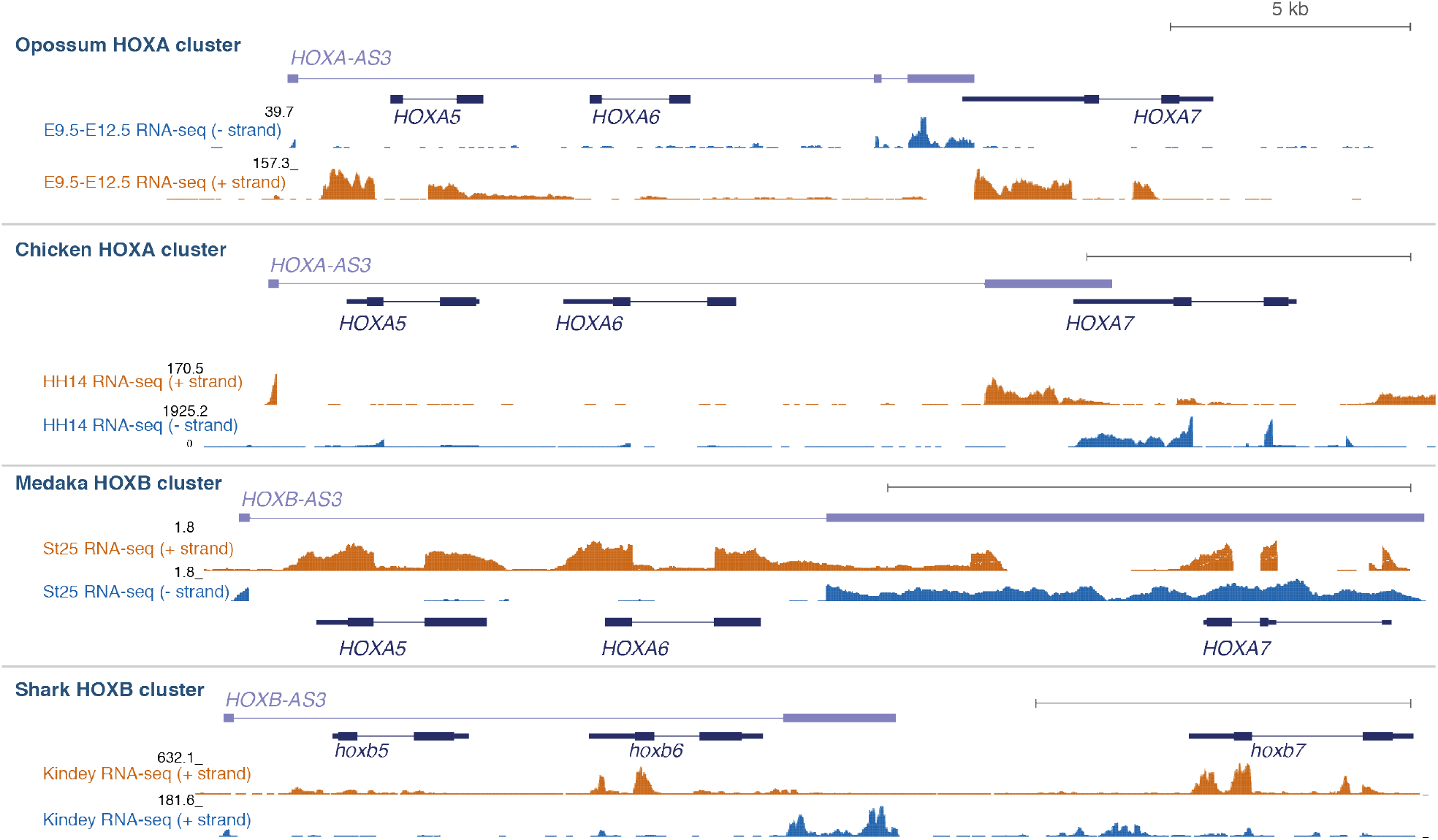
RNA-seq read coverage support for gene models of HOXA-AS3 and HOXB-AS3 homologs. In each species, annotated or reconstructed gene models are shown for *HOXA-AS3* or *HOXB-AS3* (the strand from which they are produced is defined as the ‘+’ strand) and the protein-coding HOX5–7 genes (transcribed from the ‘-’ strand). RNA-seq data are from the following datasets: SRP023152 (opossum), SRP041863 (Chicken), GSE136018 (Medaka), and SRP013772 (Shark).

**Figure S2:**
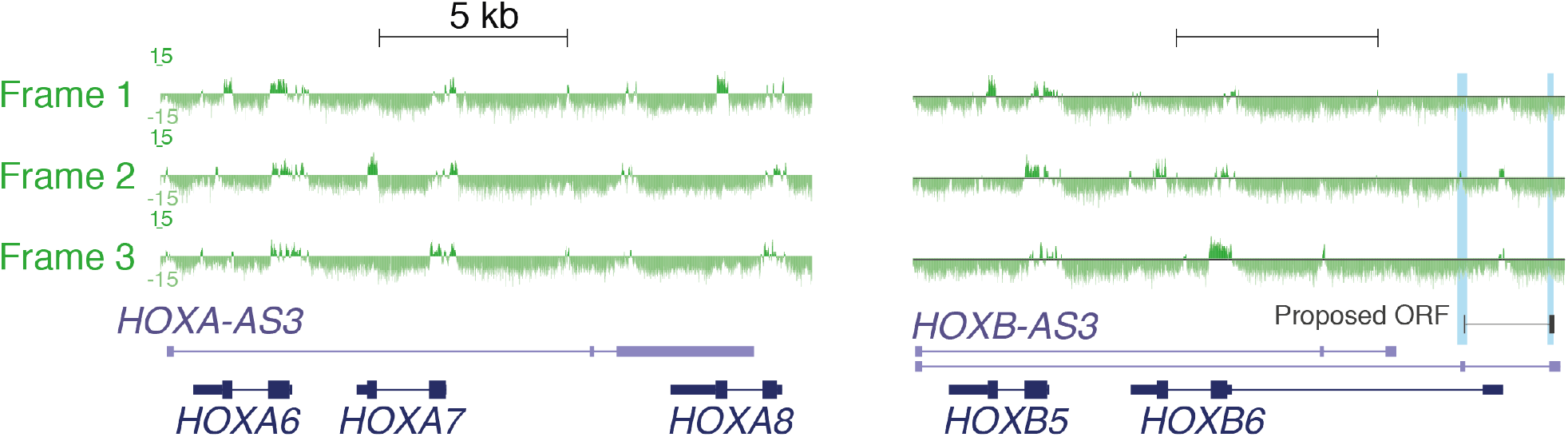
PhyloCSF scores for the HOXA-AS3 and HOXB-AS3. PhyloSCF scores (Lin et al. 2011) taken from the PhyloCSF UCSC genome browser, for each of the three frames for the ‘+’ strand from which HOXA-AS3 and HOXB-AS3 are transcribed. Position of the proposed ORF from (Huang et al. 2017) is shown.

The corresponding positions of the two lncRNAs and their high conservation in other species made us scrutinize and compare the sequences of their promoters. BLAST comparison of the corresponding promoters from HOXA and HOXB clusters found significant homology in representative vertebrate species all the way to the cartilaginous fish elephant shark (E-value 6e-31 in human, 5e-32 in mouse, 1e-21 in Xenopus, 73–80% base identity). Mapping the transcription start sites of *HOXA-AS3* and *HOXB-AS3* transcripts based on RNA-seq data (where available) suggested that the precise position of transcription initiation varies between the clusters and to a lesser extent between the species (**Fig. S3**). Among the highly conserved sequences preserved in both classes, we note a pair of tandem binding sites for the CDX1/2 proteins - CCATAAA (Verzi et al. 2010) that appear once on the sense and once on the antisense strand.

**Figure S3.**
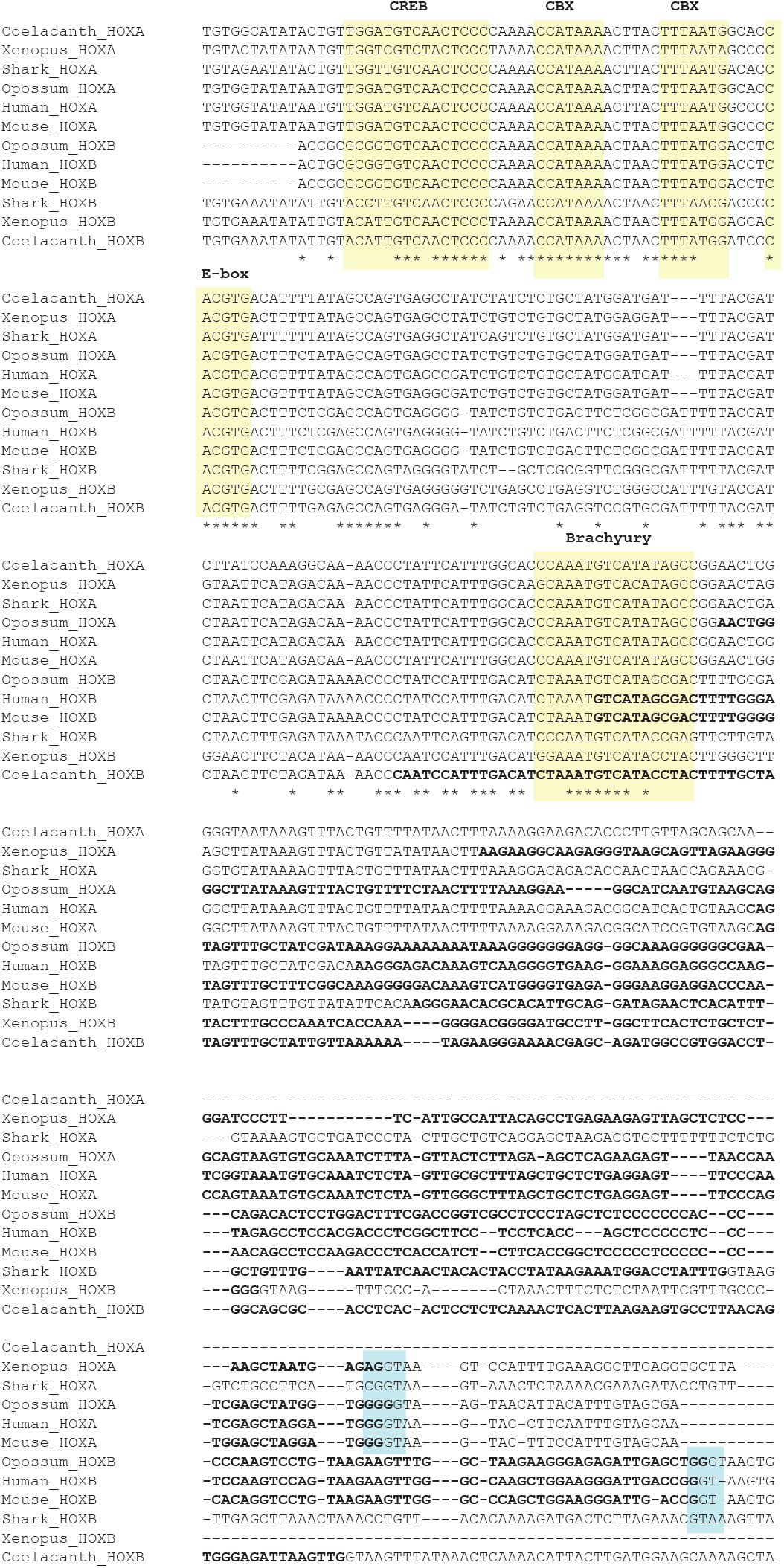
Sequence conservation and similarity in the *HOXA-AS3* and *HOXB-AS3* promoter regions. Exonic sequences (where known) are in bold. Predicted binding sites of the indicated transcription factors, taken from the UCSC genome browser are shaded in yellow. Regions of the 5’ splice sites at the end of the first exon, where known, are shaded in blue.

### *HOXA-AS3* and *HOXB-AS3* are co-expressed in embryonic and adult tissues

In order to obtain a comprehensive picture of where the two lncRNAs are expressed in both fetal and adult cell types, we relied on data from the FANTOM5.5 project (Consortium, Fantom et al. 2014), which provide strand-specific data across hundreds of cell types. Both lncRNAs were expressed in a highly specific manner and with patterns largely distinct from those of the overlapping HOX-5, HOX-6, HOX-7 genes (**Fig. 2A**). In particular, expression of *HOXA-AS3* in human and in mouse was more tissue-specific than that of other considered genes (with the exception of HOXA6 in human, **Fig. 2A**) and more closely resembled *HOXB-AS3* than any of the genes whose transcription units it overlapped. The samples in which *HOXA-AS3* and *HOXB-AS3* were expressed (**Table S1**) were mostly embryonic or derived from embryonic stem cells. In human, both lncRNAs were co-expressed during late-stage differentiation of embryonic stem cells to embryoid bodies. In mouse, both *Hoxaas3* and *Hoxb5os* were co-expressed in the E8.5 mesoderm in the neonate intestine (consistent with the single cell RNA-seq data, **Fig. 2B**).

**Figure 2.**
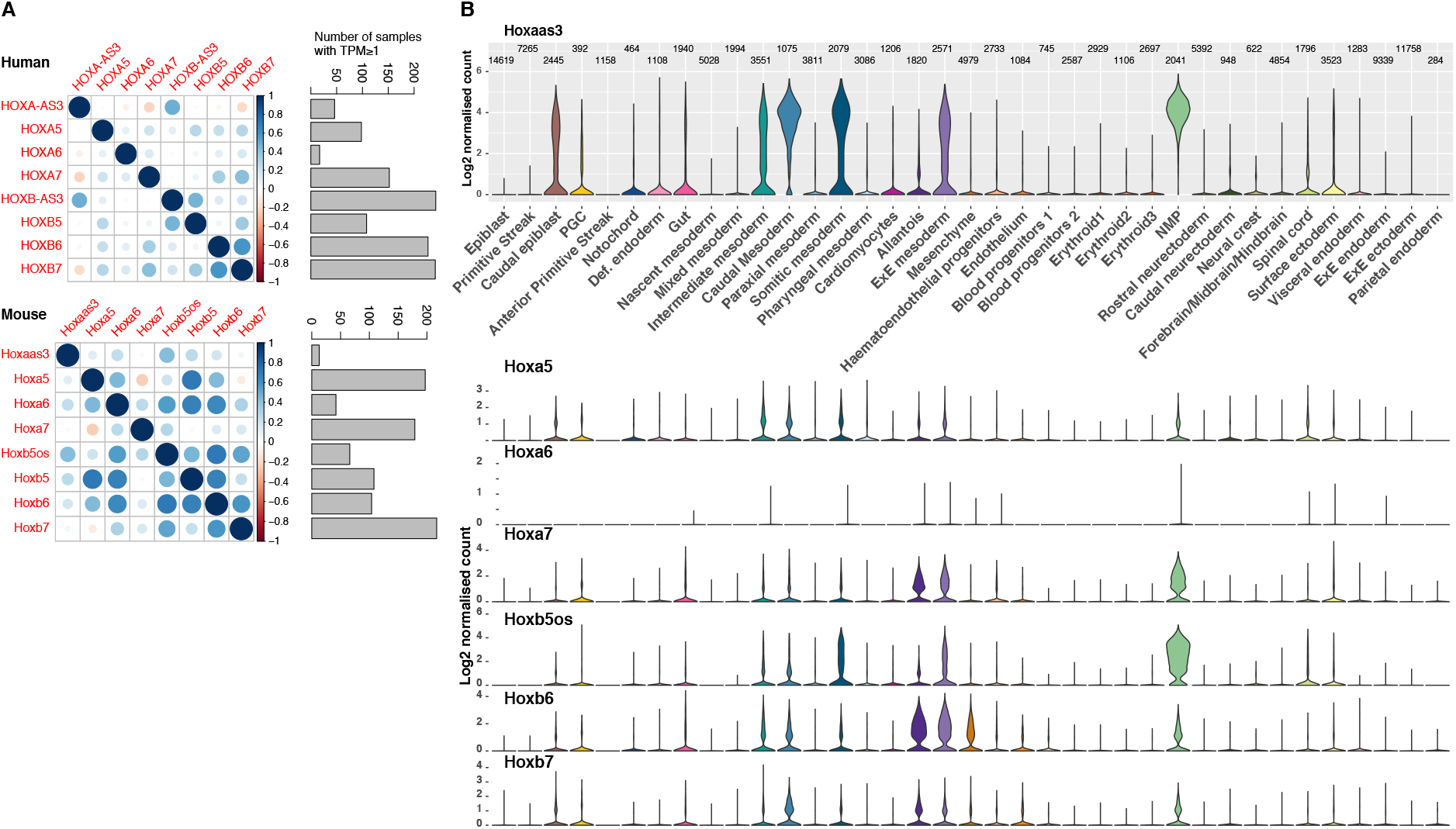
Expression of HOXA-AS3 and HOXB-AS3 in adult and embryonic tissues. **(A)** Left: Correlation coefficients between log-transformed FANTOM5 expression levels in hundreds of samples for the indicated genes. Right: Number of samples in which each gene is expressed at TPM≥1. **(B)** Expression levels of the indicated genes in clusters of single cells during gastrulation, data from (Pijuan-Sala et al. 2019).

We next examined *Hoxaas3* expression in adult mouse tissues in the Tabula Muris scRNA-seq dataset (*Hoxb5os* is not annotated in this dataset). The only cell type where there was appreciable expression were Goblet and epithelial cells from the large intestine (**Fig. S4**), consistent with our more detailed analysis (see below). *HOXA-AS3* and *HOXB-AS3* thus exhibit a very high tissue specificity in the adult tissues, similarly to other lncRNAs (Cabili et al. 2011).

**Figure S4.**
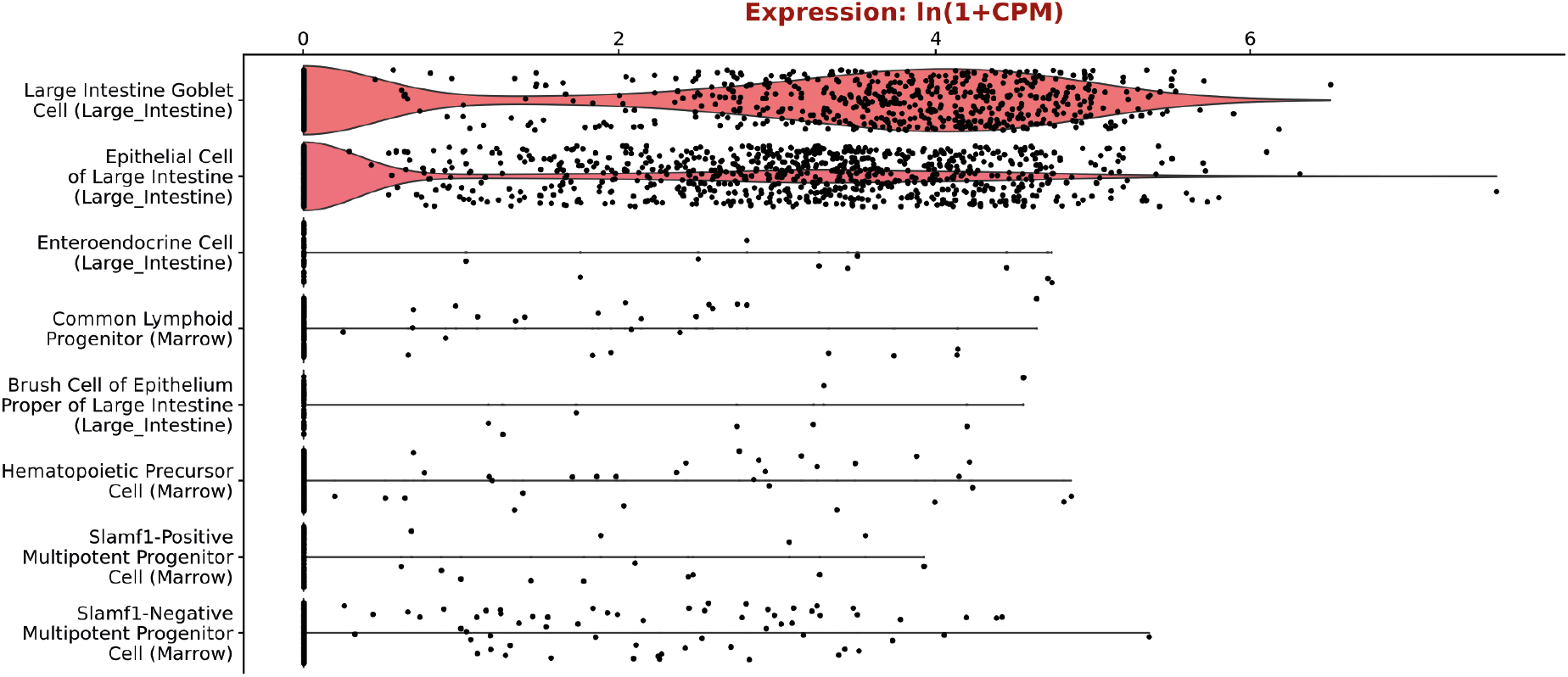
Expression of *Hoxaas3* in clusters of single cells from the Tabula Muris database (Tabula Muris Consortium et al. 2018). Eight cell groups with the highest expression are shown.

To obtain single-cell resolution on the expression of the two lncRNAs during early embryonic development, we used the large-scale single-cell dataset recently published by the Sanger institute (Pijuan-Sala et al. 2019), which profiled mouse embryos at E6.5–E8.5 (**Fig. 2B**). At these stages, *Hoxaas3* and *Hoxb5os* were generally more highly expressed than the protein-coding genes overlapping their gene bodies. As in the FANTOM data, *Hoxaas3* expression was most similar to the expression of *Hoxb5os* and the two lncRNAs were highly expressed in neuro-mesodermal progenitors (NPM in **Fig. 2B**), various mesodermal populations, caudal epiblast, and gut cells.

### *HOXA-AS3* and *HOXB-AS3* regulate their adjacent Hox genes in HT-29 cells

Inspection of ENCODE data suggested *HOXA-AS3* is not well-expressed in commonly used human cell lines, consistently with its overall low expression in adult tissues. *HOXB-AS3* is somewhat more broadly expressed, as it is expressed also in Ag04450, IMR-90, and NHLF cells. Surprisingly, there was no substantial expression of *HOXA-AS3* or *HOXB-AS3* in A549 and HUVEC cell lines where they have been previously studied (Zhu et al. 2019)(Zhang et al. 2018) (**Fig. S5** and Discussion). In contrast, ENCODE RNA-seq data showed that *HOXA-AS3* and *HOXB-AS3* are well expressed in HT-29 (**Fig. S5**) – a human colon adenocarcinoma cell line that under certain growth conditions exhibits characteristics of mature intestinal cells, such as enterocytes or mucus producing cells which have brush borders and expresses Villin and additional intestinal microvilli proteins (Martínez-Maqueda et al. 2015; Rousset 1986).

**Figure S5:**
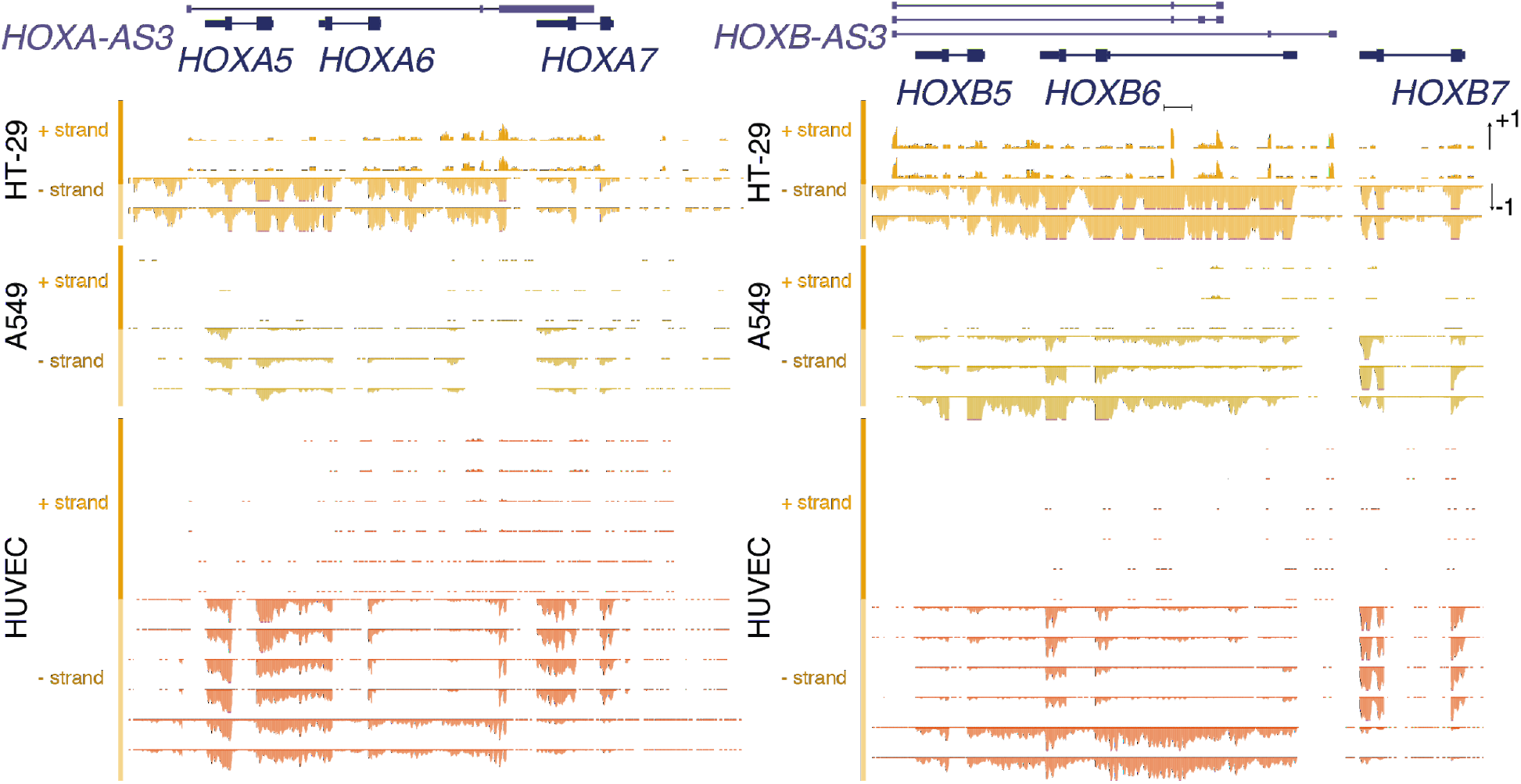
Expression of central HOXA and HOXB genes in ENCODE cell lines. Shown are selected RefSeq gene models for the indicated genes alongside RNA-seq read coverage from the indicated cell lines from ENCODE datasets of total (HT-29) or polyA-selected (A459, HUVEC) RNA on the indicated strand. *HOXA-AS3* and *HOXB-AS3* are transcribed from the ‘+’ strand and the protein-coding genes from the ‘-’ strand.

In order to perturb the expression of *HOXA-AS3* and *HOXB-AS3*, we first used CRISPR interference (CRISPRi) (Gilbert et al. 2013) – a catalytically inactive version of Cas9 (dCAS9) optionally fused to a KRAB domain (dCas9-KRAB) together with guide RNAs (gRNAs) directed to a region downstream to the TSS of the target (Qi et al. 2013). We transfected HT-29 cells with pools of three gRNAs targeting *HOXA-AS3* and *HOXB-AS3* promoters and dCAS9-KRAB vectors, which reduced lncRNA levels by 50%–80% compared to cells transfected with the dCas9-KRAB vector and an empty gRNA plasmid (**Fig. 3A-B**). As *HOXA-AS3* levels are reduced, *HOXA5, HOXA6* and *HOXA7* RNA levels are also down-regulated by 30–40% (**Fig. 3A**). Similarly, *HOXB-AS3* knockdown (KD) was followed by a down regulation of *HOXB5* and *HOXB6* by 50–70% (**Fig. 3B**).

**Figure 3.**
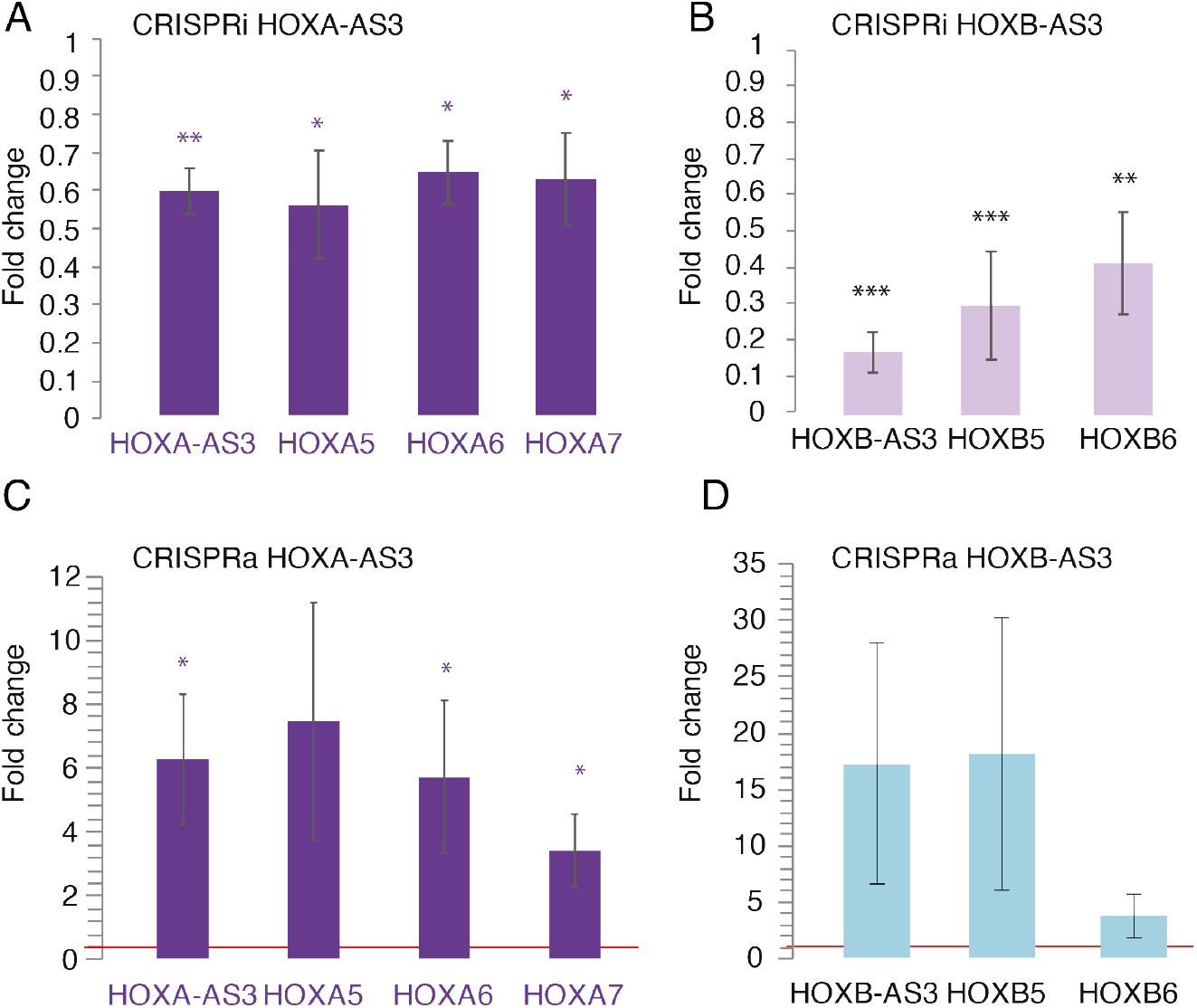
CRISPR inhibition and activation of *HOXA-AS3* and *HOXB-AS3* in HT-29 cells. **(A)** Changes in expression of the indicated genes is shown following inhibition of *HOXA-AS3*. n=4. (**B**) As in A, following inhibition of *HOXB-AS3*. n=4. (**C**) As in A, following activation of *HOXA-AS3*. n=4 (**D**) As in A following activation of *HOXB-AS3*. Normalized to actin. Two-sided t-test. * - P<0.05, ** - P<0.005, *** - P<0.0005. Two-sided t-test compared to the transfection control.

Next, we tested the effect of over-expression (OE) of the lncRNAs using CRISPR activation (CRISPRa) – dCas9 fused to VP64 transcriptional activation domain and directed by the sgRNAs to a region upstream to the *HOXA-AS3* and *HOXB-AS3* TSSs. In this system, the VP64 domain recruits the transcription machinery to activate expression of the lncRNA of interest (Qi et al. 2013). OE in HT-29 cells resulted in effects opposite to those observed following lncRNA KD, as it increased expression of the adjacent genes (**Fig. 3C-D**). For *HOXA-AS3* we also tested activation of the promoter in a MCF-7 cell line that does not normally express it or the neighboring genes and saw an upregulation of the adjacent genes as well (**Fig. S6**).

These results suggest that *HOXA-AS3* and *HOXB-AS3* production or their RNA products have a positive regulatory effect on the expression of the neighboring genes HOX5–7 genes.

**Figure S6.**
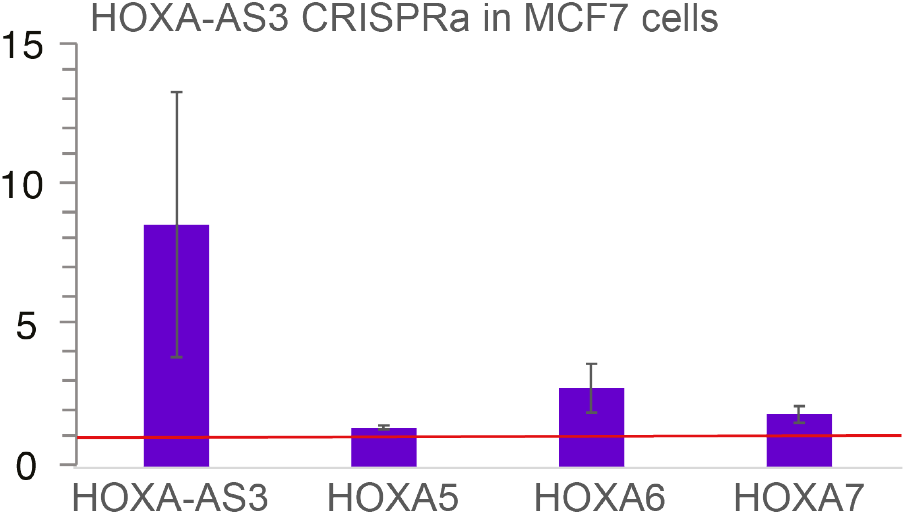
CRISPR activation of *HOXA-AS3* in MCF7 cells. Changes in expression of the indicated genes is shown following activation of *HOXA-AS3*. n=2.

### *HOXA-AS3* and HOXB-AS3 RNA products are required for their *cis*-regulatory activity

lncRNAs can act in *cis* or in *trans* to carry out their functions. In order to differentiate between the potential effects on chromatin caused by the use of the KRAB effectors and the transcription or the RNA products of *HOXA-AS3* and *HOXB-AS3*, we used RNAi to target the RNA products of *HOXA-AS3* and *HOXB-AS3*. First we transfected siRNA pools targeting *HOXA-AS3* or *HOXB-AS3* into HT-29 cells. This resulted in a substantial reduction in RNA levels for both *HOXA-AS3* and *HOXB-AS3* and a concomitant reduction in the expression of neighboring genes that was similar to the effects observed with CRISPRi (**Fig. 4A-B**). When *HOXA-AS3* was reduced by 60%, *HOXA5*/*6*/*7* were significantly downregulated by 20-45% (**Fig. 4A**). Similarly, when *HOXB-AS3* was reduced by ~40%, there was a significant downregulation of *HOXB5* and *HOXB6* (**Fig. 4B**). As an alternative approach, a stably expressed shRNA targeting *HOXA-AS3* introduced via a lentiviral infection led to a stronger effect with the same trend as that observed using CRISPRi and siRNA, where KD of the lncRNA was accompanied by a decrease of expression of the neighboring genes (**Fig. 4C**). *HOXA-AS3* and *HOXB-AS3* RNA products are therefore important for regulation of the adjacent genes.

**Figure 4.**
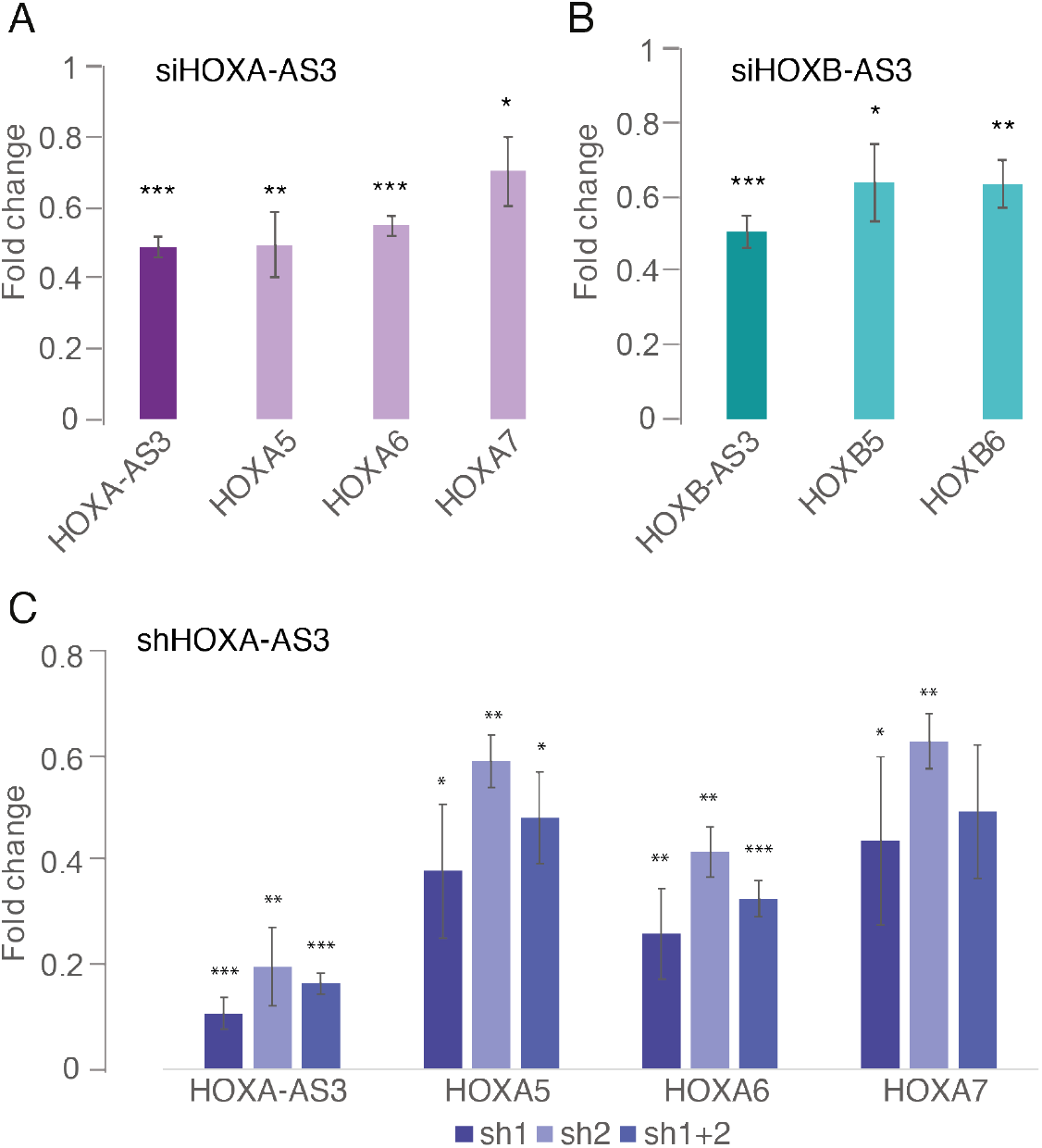
RNA products of *HOXA-AS3* and *HOXB-AS3* are required for regulation of their adjacent Hox genes. qRT-PCR measurements of the indicated genes in HT-29 cells treated with the indicated reagents. Normalized to Actin. n=4 for siHOXA-AS3 and siHOXB-AS3. n=3 for shHOXA-AS3. * - P<0.05, ** - P<0.005, *** - P<0.0005. Two-sided t-test compared to the transfection control.

### *HOXA-AS3* is localized in the both the nucleus and cytoplasm of HT-29 cells

We next focused on *HOXA-AS3* and characterized its precise expression pattern at higher resolution, as it is more narrowly expressed compared to *HOXB-AS3*, and also has a longer exonic sequence which permits the use of Stellaris smFISH protocol with 96 exonic probes for the human *HOXA-AS3* and 94 for the mouse *Hoxaas3* (**Table S2**), whereas only 24 probes were possible for *HOXB-AS3*.

We first analyzed the subcellular localization of *HOXA-AS3* and *HOXA5* in HT-29 cells (**Fig. 5A**). We observed variable expression of both genes among cells, in some of the cells we could detect expression of only one of the transcripts, while others expressed both genes. *HOXA-AS3* transcript was detectable in just ~15% of the >100 imaged cells, in up to 3 foci per cell and with localization mainly in the nucleus, though it could also be detected in the cytoplasm.

**Figure 5:**
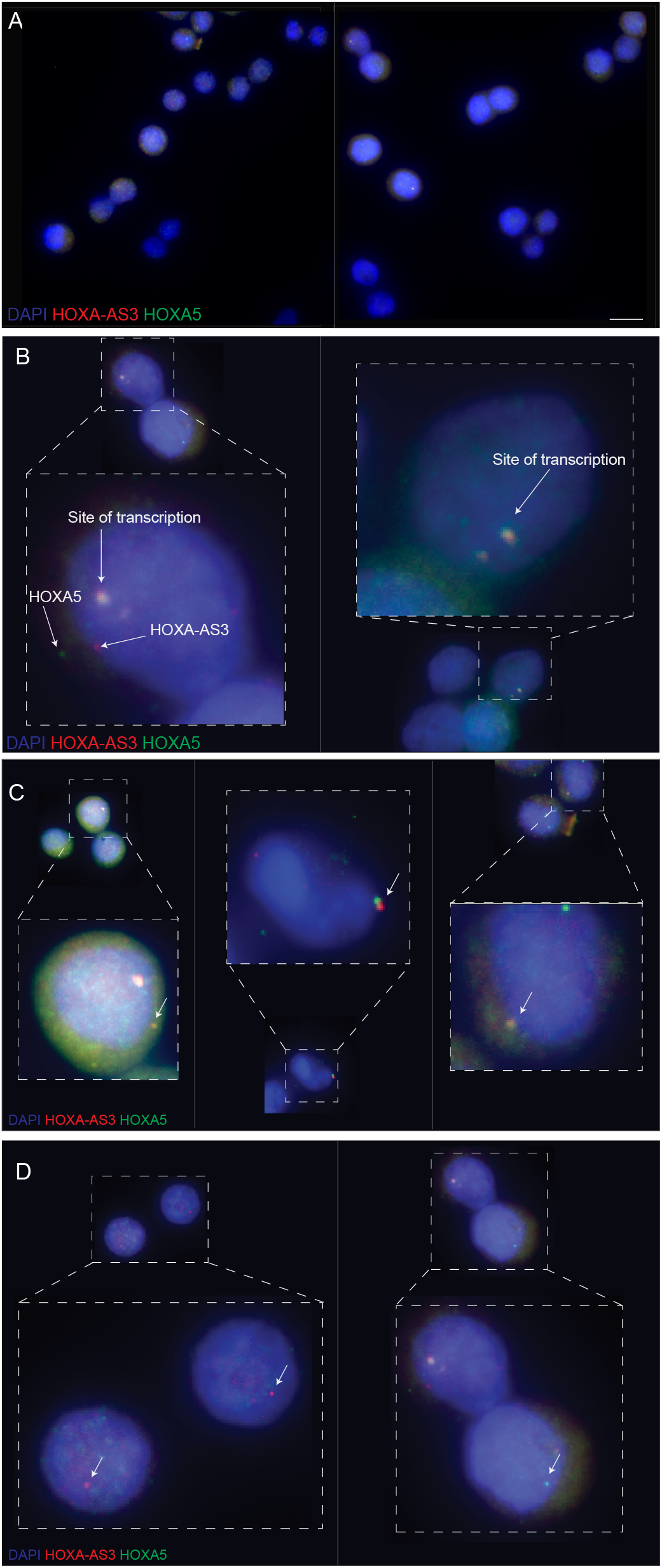
Single-molecule FISH detection of *HOXA-AS3* and *HOXA5* in HT-29 cells. **(A)** *HOXA-AS3* (red) and *HOXA5* (green) transcripts in a sample of HT29 cells. Scale bar: 10 µm. **(B)** *HOXA-AS3* and *HOXA5* are co-localized at their presumed site of transcription. **(C)** *HOXA-AS3* and *HOXA5* are occasionally co-localized in the perinuclear area (white arrow). **(D)** *HOXA-AS3* and *HOXA5* are occasionally expressed separately.

Interestingly, in some of the cells that express both *HOXA-AS3* and *HOXA5* we detected a rare yet highly specific co-localization in the perinuclear area (**Fig. 5B**). As expected from their genomic co-location, *HOXA-AS3* and *HOXA5* are co-localized in what is likely their site of transcription in the nucleus (**Fig. 5B**).

### *HOXA-AS3* is expressed in a specific subset of colon epithelial cells

As HT-29 cells contain a mixture of cellular states from the colon epithelium (Rousset 1986; Martínez-Maqueda et al. 2015), *HOXA-AS3* expression in a small subset of cells may imply that it is only found in a defined subpopulation of cells. We therefore analyzed the expression pattern of *HOXA-AS3* and *Hoxaas3* in normal intestinal epithelial cells, using single-cell RNA sequencing (scRNA-seq) data.

In scRNA-seq data from the human colon scRNA-seq data, HOXA-AS3 was expressed predominantly in epithelial cells, and within those it was detected specifically in tuft and immature goblet cells, that are deep crypt goblet cells that are part of the stem cell niche (Clevers 2013) (**Fig. 6A**). Similarly, in the mouse small intestine (Haber et al. 2017) *HOXA-AS3* is mainly expressed in tuft cells at comparable expression levels to the tuft marker *Dclk1* (**Fig. 6B**). In contrast, in the mouse colon scRNA-seq *Hoxaas3* is mainly detected in goblet cells (**Fig. 6C**). In order to examine expression in intact tissue, we performed smFISH for *Hoxaas3* in the jejunum of the mouse small intestine, which contains a relatively high fraction of goblet cells, and compared it to smFISH of the goblet cell marker *Gob5*, the tuft cell marker *Dclk1*, and *Atoh1* marking intestinal secretory precursor cells, including immature goblet and tuft cells. We observed that *Hoxaas3* was enriched in the early immature goblets and in the secretory precursor cells (**Fig. 6D**).

**Figure 6.**
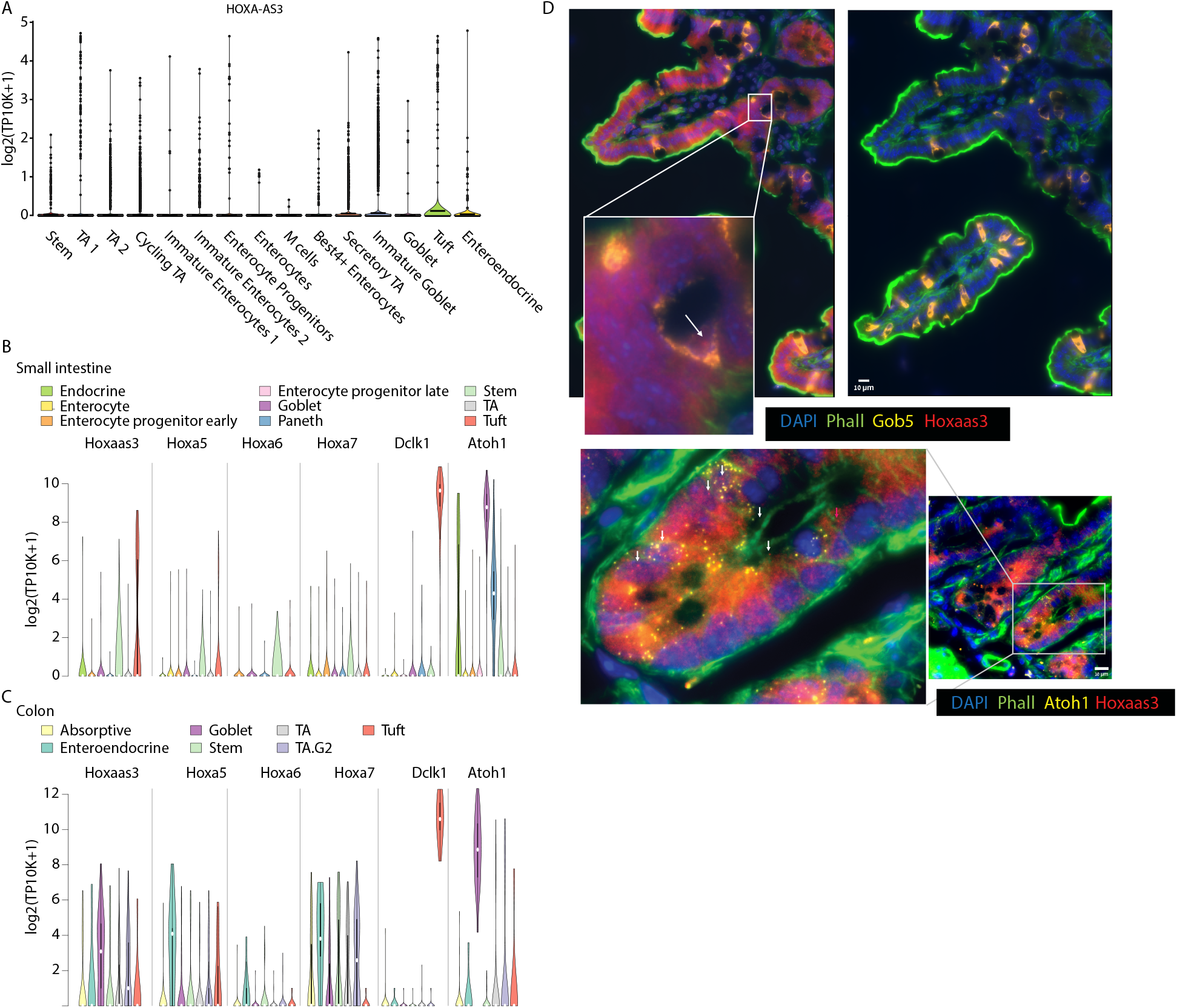
Expression of HOXA-AS3 in the human and mouse gut. **(A)** Expression of HOXA-AS3 in single cells of the human colon (data from (Smillie et al. 2019)). **(B-C)** Expression of the indicated genes in scRNA-seq from the mouse small intestine (**B**) and colon (**C**). Data from (Haber et al. 2017). **(D)** smFISH of HOXA-AS3, *GOB5* and *Atoh1* expression in mouse intestine. Scale bar:10µm

scRNA-seq and smFISH from both human and mouse samples thus supports the notion that *HOXA-AS3* is expressed in a specific subpopulation, which may explain the apparently variable expression pattern that we observed in HT-29.

### *HOXA-AS3* and *HOXB-AS3* are induced during early differentiation of human embryonic stem cells towards endoderm

As both *HOXA-AS3* and *HOXB-AS3* were more highly expressed in embryonic stages compared to adult tissues, we next wanted to evaluate the expression and activities of *HOXA-AS3* and *HOXB-AS3* during early developmental transitions. Endoderm is one of the three primary germ cell layers, and endoderm patterning is controlled by a series of reciprocal interactions with nearby mesoderm tissues. As development proceeds, broad gene expression patterns within the foregut, midgut, and hindgut become progressively refined into precise domains from which specific organs will arise. Human embryonic stem cells can be differentiated towards endodermal cell lineages in a robust manner, resulting within seven days in three different populations – anterior foregut (AFG), posterior foregut (PFG) and midgut/hindgut (MHG), using a protocol established by Loh et al. (Loh et al. 2014) **(Fig. S7A-B**). During this differentiation process a graded, spatially collinear Hox gene expression is observed, after *in-vitro* patterning, whereby PFG cells express 3’ anterior Hox genes (e.g. *HOXA1*) and MHG cells express 5’ posterior Hox genes (including *HOXA10*) (Loh et al. 2014)

**Figure S7:**
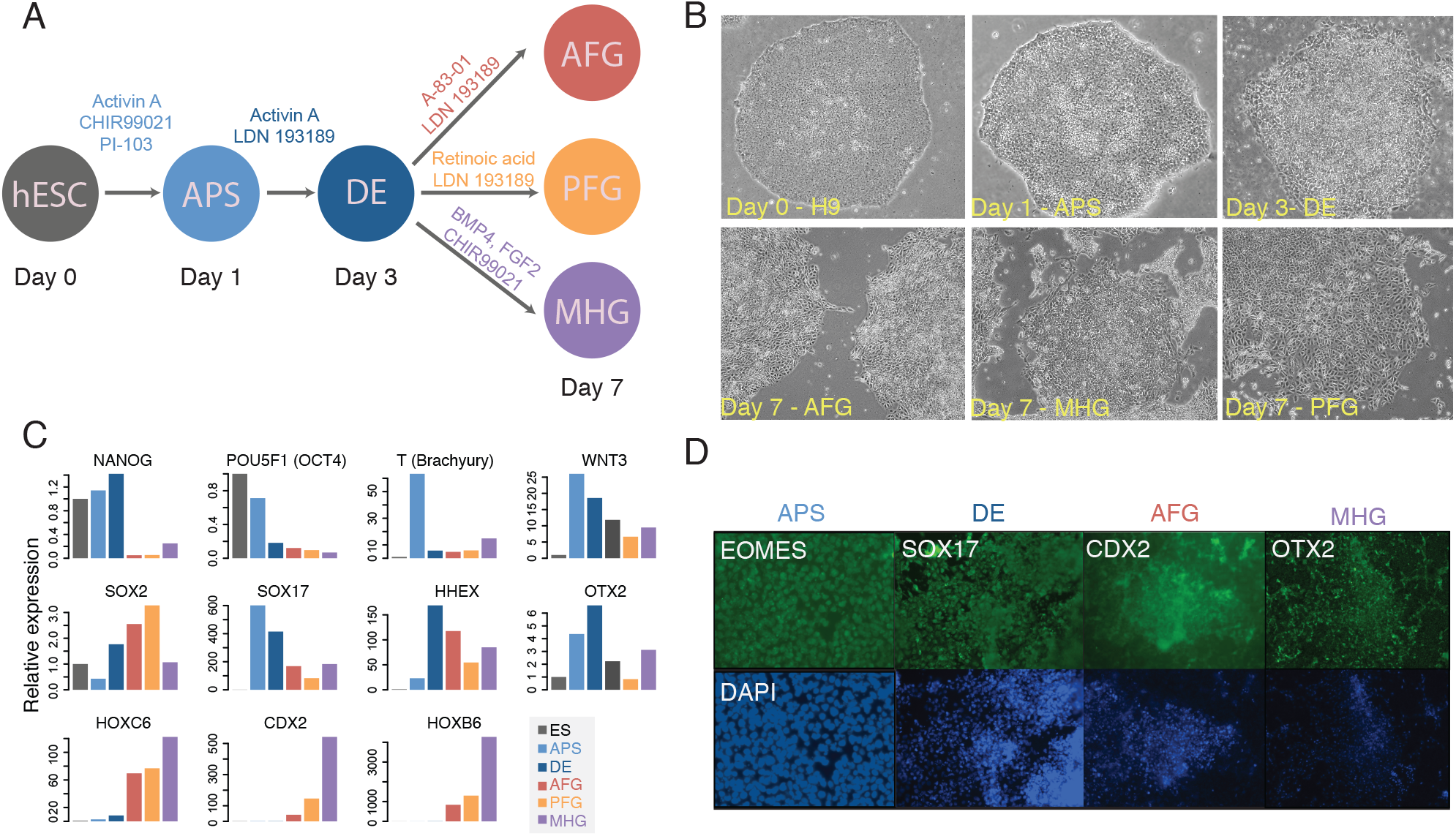
hESC endodermal differentiation overview. **(A)** Endodermal differentiation process and signaling molecules. **(B)** Characterization of different cell morphology in different stages of the differentiation. **(C)** RNA levels and expression dynamics were measured by qRT-PCR at different stages of endodermal differentiation, and normalized to actin. **(D)** Immunofluorescence stainings of human ESCs differentiated in different stages of differentiation.

Pluripotent hESCs and cells from each stage of the differentiation were validated by multiple markers (**Table S3**) using qRT-PCR (**Fig. S7C**) and by immunostaining (**Fig. S7D**), matching the expression patterns observed in the RNA-seq data from (Loh et al. 2014) (**Fig. 7A**), *HOXA-AS3* and *HOXB-AS3* were strongly induced and expressed only in the MHG population, alongside their adjacent HOX-6 and HOX-7 genes, whereas *HOXA5* and *HOXB5* were alse expressed in PFG cells (**Fig. 7A**).

**Figure 7.**
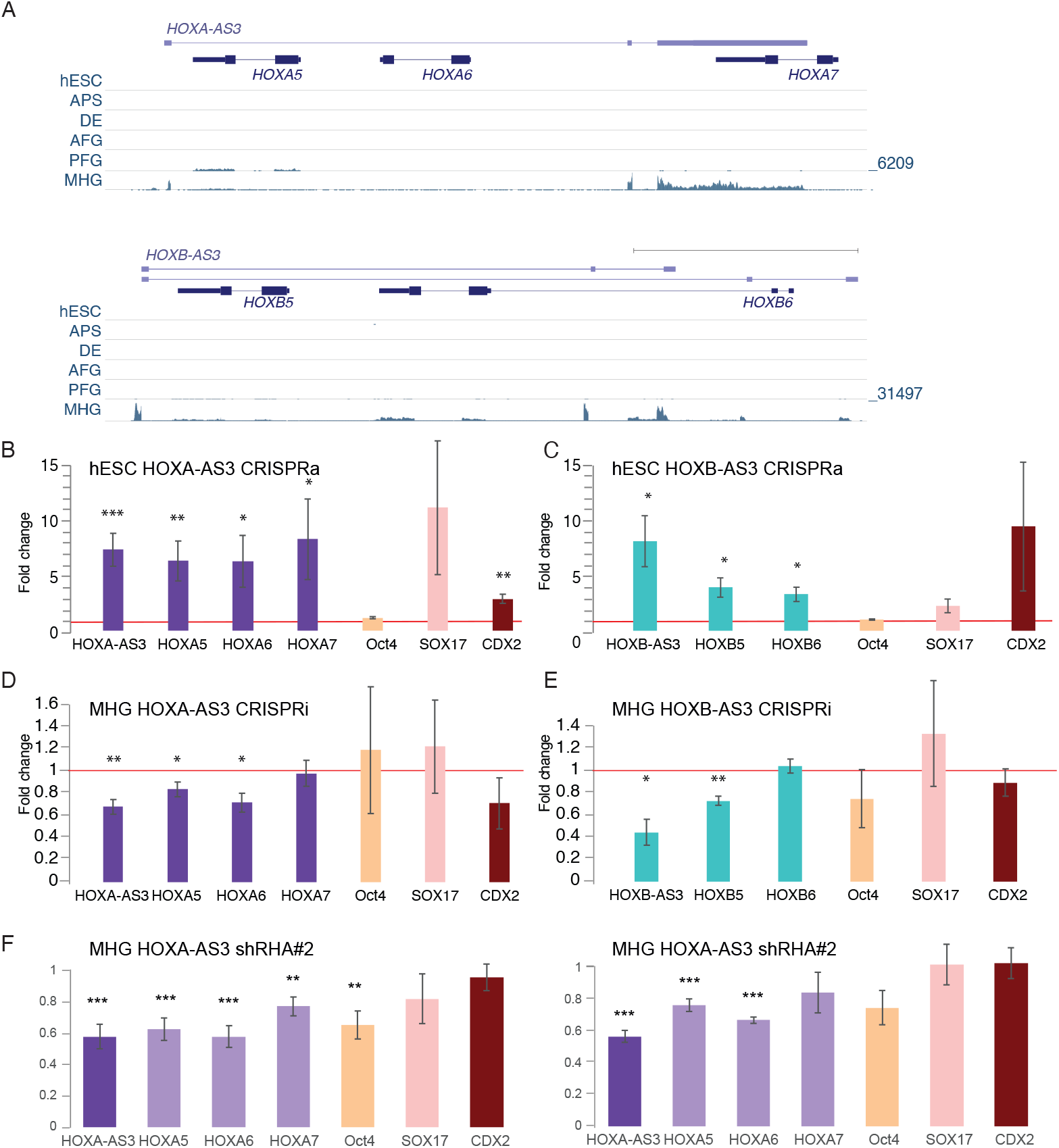
Function of HOXA-AS3 and HOXB-AS3 during endodermal differentiation of hESCs. **(A)** Read coverage in RNA-seq data from (Loh et al. 2014) for shown parts of the HOXA (top) and HOXB (bottom) clusters. In each cluster all the tracks are normalized together. **(B-C)** Expression levels estimated by qRT-PCR for the indicated genes in hESCs following CRISPRa-mediated 48h activation of *HOXA-AS3 (n=7/4)* (**B**) and *HOXB-AS3* (n=3) (**C**). (**D-E**). Expression levels estimated by qRT-PCR in MHG cells following CRISPRi-mediated repression of *HOXA-AS3 (n=3)* (**D**) and *HOXB-AS3* (n=3) (**E**). **(F)** Changes in expression of the indicated genes following infection of hESCs with two separate HOXA-AS3 shRNAs, followed by differentiation to MHG. n=6. * - P<0.05; ** - P<0.005; *** - P<0.005. Two sided t-test. Errors bars - SEM.

### *HOXA-AS3* and *HOXB-AS3* regulate expression of their adjacent Hox genes during hESC differentiation

To study the functions of *HOXA-AS3* and *HOXB-AS3* during early steps of stem cell differentiation, we established dCas9-expressing H9 hESCs, using viral infection of Tet-dependent inducible versions of the dCAS9 and dCas9-VP64. We preferred to avoid the use of dCas9-KRAB in this system, as we were able to obtain efficient KD using dCas9 alone, which does not by itself directly affect chromatin modifications. We then established derivatives of these stable lines expressing specific gRNAs targeting the promoters of *HOXA-AS3* or *HOXB-AS3*.

After 48 hr of doxycycline (Dox) addition to the dCas9-VP64 expressing lines, we observed an up-regulation of *HOXA-AS3* and *HOXB-AS3* lncRNAs in their respective lines (**Fig. 7B-C**). Furthermore, we observed up-regulation of the genes adjacent to these lncRNAs, even though neither of these genes are normally expressed in hESCs, and the chromatin of the HOXA cluster in hESC is in an inactive conformation (Soshnikova and Duboule 2009, 2008; Kashyap et al. 2011; Varlakhanova et al. 2011). Activation of *HOXA-AS3* in hESCs resulted in increased expression of *HOXA5*-*7* (**Fig. 7B**). Similarly, *HOXB-AS3* activation led to an activation of *HOXB5* and *HOXB6* (**Fig. 7C**). Next, we tested for changes in the pluripotency and differentiation markers in the CRISPRa lines. Although there was no remarkable change in the Oct4 pluripotency marker, we observed an increase in endodermal markers – *HOXA-AS3* or HOXB-AS3 overexpression (OE) lines led to an upregulation of *Sox17*, a definitive endoderm marker known to be required for normal development of the definitive gut endoderm (Kanai-Azuma et al. 2002) and in *Cdx2* levels, a marker of later stages of endodermal differentiation, expressed mainly in the MHG cells (**Fig. 7C**).

Next we wanted to examine the effect of reducing the levels of *HOXA-AS3* and *HOXB-AS3* during endodermal differentiation at time points at which they are endogenously induced during the third stage of differentiation, as the cells are transitioning from DE to MHG (**Fig. 7A**). For both *HOXA-AS3* and *HOXB-AS3* we obtained a ~50% KD using the Dox-inducible dCAS9, and targeting *HOXA-AS3* resulted in downregulation of *HOXA5*/*6*/*7* (**Fig. 7D**), with a relatively smaller effect on HOXA7, which is generally expressed at low levels in MHG cells. KD of *HOXB-AS3* led to downregulation of *HOXB5* (**Fig. 7E**). In both cases we observed no major changes in expression of markers for pluripotency (*Oct4*), endoderm (*Sox17*) and mid/hindgut (*Cdx2*) (**Fig. 7D-E**).

In order to study the role of *HOXA-AS3* RNA product during hESC differentiation, we used the two shRNA constructs described above to generate hESC lines where *HOXA-AS3* is stably targeted by RNAi. In this system, a stable reduction of *HOXA-AS3* expression also has a similar effect on HOXA5–7 (**Fig. 7E**). There was also a significant reduction in levels of Oct4, although its expression is low at the MHG stages, and so the physiological significance of this reduction is unclear.

### CDX transcription factors regulate expression of *HOXA-AS3* and *HOXB-AS3*

Expression domain of *HOXA-AS3* closely overlaps the known expression domains of the *CDX2* transcription factor (TF) in the intestine, which is expressed mainly in the maturing cells that are closer to the crypt (Grainger et al. 2010). This similar pattern of expression in intestinal epithelium, combined with the pair of CDX1/2 binding sites found in the ultra-conserved part of the *HOXA-AS3* and *HOXB-AS3* promoters (**Fig. S3**), and the ChIP-seq data from mouse epithelium (Kumar et al. 2019) (**Fig. S8A**) suggest that CDX1/2 TFs may activate *HOXA-AS3* and *HOXB-AS3*. To test this we used siRNAs to KD CDX1/2 in HT-29 cells. This resulted in a 40–75% reduction in CDX1/2 expression and a significant reduction of 30–70% in *HOXA-AS3* leves, while *HOXB-AS3* showed a significant reduction of 60% only upon CDX1 KD (**Fig. S8B**). This finding, together with the smFISH and scRNA-seq data, supports the notion that CDX TFs drive *HOXA-AS3* (and to a possibly lesser extent also *HOXB-AS3*) expression in a subpopulation of endodermal cells, enabling them in turn to function as regulators of their neighboring Hox genes.

**Figure S8:**
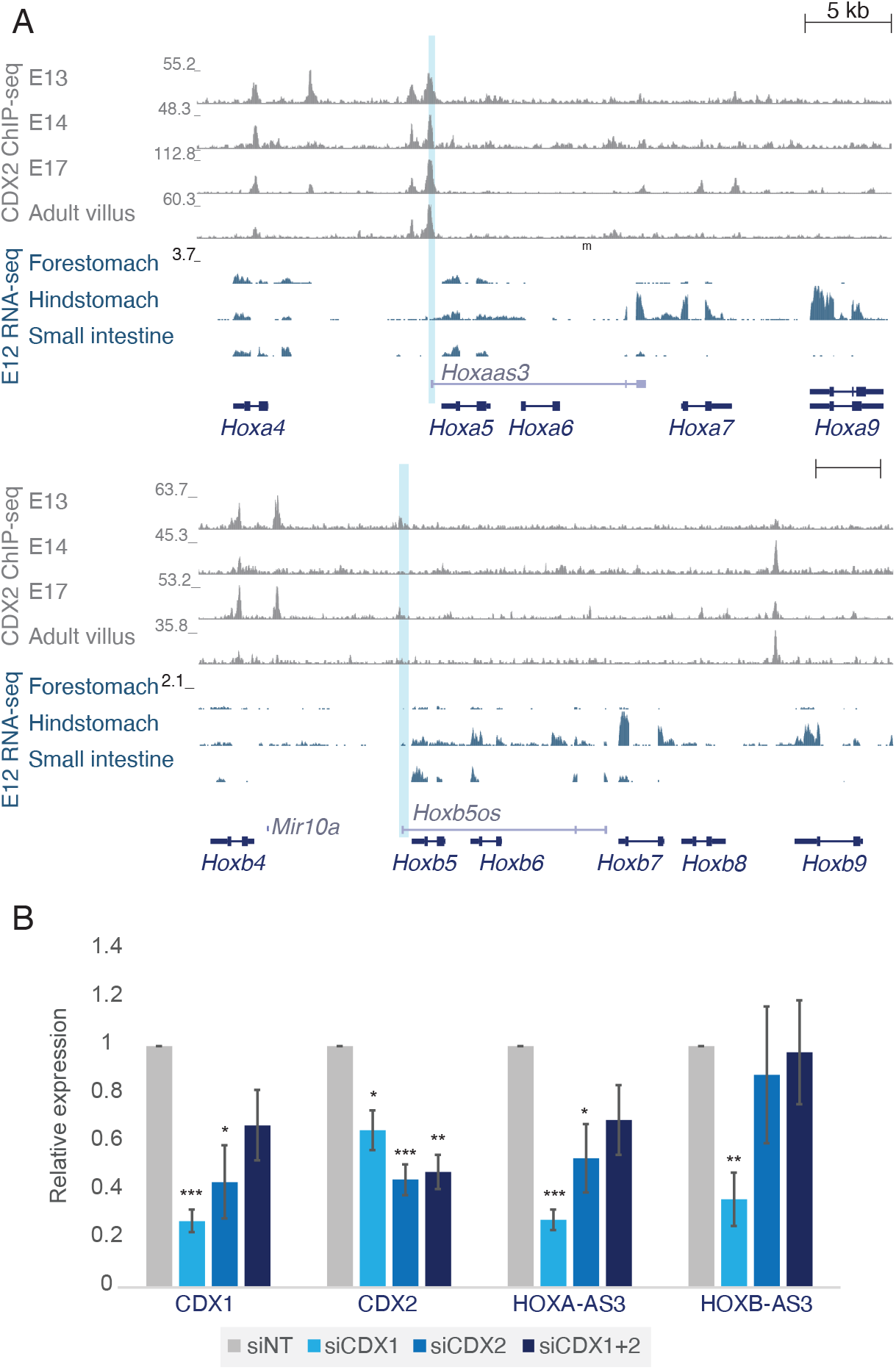
Regulation of *HOXA-AS3* and *HOXB-AS3* by CDX1/2 transcription factors. CDX2 ChIP-seq read coverage from the gut at the indicated stage and RNA-seq at E12 in the indicated tissue, data from (Kumar et al. 2019). The region corresponding to the promoters of *Hoxaas3* and *Hoxb5os* is shaded. **(B)** qRT-PCR of the indicated genes in HT-29 cells transfected with the indicated siRNAs. Normalized to siNT and to actin. n=3 * - P<0.05, ** - P<0.005, *** - P<0.0005. Two-sided t-test.

## Discussion

We found here that *HOXA-AS3* and *HOXB-AS3* are ultraconserved lncRNAs which demonstrate high conservation in promoter sequence, genomic configuration and regulation that underpin similar expression patterns in specific biological processes, and relate to related functions in regulating expression of their proximal genes. The expression of *HOXA-AS3* and *HOXB-AS3* during embryonic development is particularly high in intestine-specifying lineages such as the MHG cell population that emerges during endodermal differentiation of hESCs, in hindstomach and small intestine epithelial cells in E12 during mouse development (**Fig. S8**). Moreover we observe co-expression of *HOXA-AS3* and *HOXB-AS3* in the adult intestine and colon in human and in mouse, specifically in cells that transition from the stem cell niche to fully specified intestinal cells in the crypt, presumably utilizing some of the same regulatory programs that are used during early development. smFISH in mouse intestine showed specific enrichment of HOXA-AS3 expression in early immature goblet cells and in the secretory precursor cells, highlighting the expression timing to mid-differentiation – the phase where the cells are committing and acquiring their specific fate, in concurrence to its induction in hESC differentiation, and potentially related to expression in only a small subset of HT-29 cells in culture.

There have been several recent reports about the functions of *HOXA-AS3* and *HOXB-AS3* in other systems. *HOXA-AS3* was reported to be induced during adipogenic induction of human mesenchymal stem cells (MSCs), and its silencing promoted proliferation of MSCs and inhibited osteogenesis *in vitro* and *in vivo*, in both human and mouse cells (Zhu et al. 2016). Positive effects of *HOXA-AS3* on proliferation and migration *in vitro* and during tumorigenesis *in vivo* were also observed in glioma cells (Wu et al. 2017). Another study found that *HOXA-AS3* promoted proliferation, migration and invasion in A549 lung carcinoma cell line and tumor growth *in vivo* (Zhang et al. 2018), where it was found in both the nucleus and the cytoplasm, consistent with our data in HT-29 cells. In that study *HOXA-AS3* was suggested to positively regulate of *HOXA6*, as siRNA-mediated KD of *HOXA-AS3* reduced levels of *HOXA6* mRNA and protein (but not those of *HOXA5*) in A549 cells. A more recent publication extended the positive effects of *HOXA-AS3* on proliferation to additional non-small-cell lung carcinoma cell lines (Lin et al. 2019). These studies are overall consistent with our observations that in normal tissues *HOXA-AS3* is preferentially expressed in proliferating progenitors. Lin et al. found *HOXA-AS3* to have a negative effect on *HOXA3* expression, by binding to both *HOXA3* mRNA and *HOXA3* protein. Lastly, HOXA-AS3 was recently proposed to regulate NF-kappaB signalling in HUVECs (Zhu et al. 2019). Notably, in ENCODE RNA-seq data *HOXA-AS3* is undetectable in both A549 cells and HUVECs (which do express *HOXA6* and *HOXA7*), whereas it is well-expressed in the HT-29 cells we used in this study (**Fig. S5**).

*HOXB-AS3* was reported to be down-regulated in colorectal cancers and to produce a 53 aa protein conserved in primates (Huang et al. 2017). Notably, PhyloCSF scores throughout *HOXB-AS3* are negative (**Fig. S1**), so it is very unlikely that it encodes a conserved protein. In colorectal cancer cells *HOXB-AS3* was shown to inhibit cell proliferation (Huang et al. 2017). In NPM1-mutated acute myeloid leukemia cells, *HOXB-AS3* does not associate with polysomes and promotes cell proliferation in both human and mouse leukemia cells (Huang et al. 2019)(Papaioannou et al. 2019). Interestingly, in this system, KD of *HOXB-AS3* using antisense oligonucleotides did not affect expression of other HOX genes, but rather regulated expression of ribosomal RNA, in *trans*, via interaction with EBP1 (Papaioannou et al. 2019). There is therefore evidence of trans-acting activities of *HOXB-AS3*. We note that our findings about cis-acting regulation of *HOXB6* and *HOXB7* by *HOXB-AS3* do not exclude these additional functions and in fact it is likely that lncRNAs that are robustly expressed and highly conserved have aquired additional, species-or clade-specific functions during evolution.

Various mechanism *for* cis-acting regulation of gene expression by lncRNAs have been demostrated in different systems (Gil and Ulitsky 2020). Future studies will elucidate the mechanism underlying the regulation of HOX5–7 gene expression by *HOXA-AS3* and *HOXB-AS3*, which may resemble those of other lncRNAs. Genome editing of the loci can be particularly powerful for promoting understanding of lncRNA biology, but it is particularly difficult to perform and interpret in the Hox clusters, due to the high density of gene regulatory elements within the clusters and the complex relations between them. The most relevant systems to perform editing of the human *HOXA-AS3* and *HOXB-AS3* is likely hESCs, which can then be differentiated to MHG cells, but CRISPR-mediated editing in hESCs is inefficient (Ihry et al. 2018). Indeed, despite screening hundreds of clones, we were so far unsuccessful in obtaining homozygous deletions of the HOXA-AS3 promoter in hESCs. Mouse models carrying specific manipulations, such as insertion of polyA sites, will also be highly informative.

Some of the protein-coding genes and miRNAs in the Hox clusters were shown to be functionally equivalent to each other and to contribute differentially to organismal function via their divergent expression patterns (Greer et al. 2000; Wong et al. 2015). These orthologs formed by duplication during the formation of the four vertebrate Hox gene clusters. The paucity of known lncRNA paralogs present in different Hox clusters can be rather easily explained by the overall high rate of lncRNA evolution (Ulitsky 2016), which likely rewired the sequences and exon-intron architectures of Hox lncRNAs extensively over the past 500 million years. The numerous features we identified as shared between *HOXA-AS3* and *HOXB-AS3* suggest that at least some lncRNAs were duplicated and maintained regulatory functions in the Hox cluster throughout vertebrate evolution, during which individual clusters also acquired additional lncRNAs, some of which are functional, and that further sculpted gene expression within each cluster. Importantly, there is also evidence of extensive cross-regulation between the clusters, including by lncRNAs (Rinn et al. 2007). Future studies will examine the potential contribution of *HOXA-AS3* and *HOXB-AS3* lncRNAs to cross-cluster regulation, as well as the extent of similarity that they maintained in their modes of action.

## Materials and Methods

### Tissue culture

H9 hESC were routinely cultured on MEFs in hESC medium consisting of 500 ml of KNOCK-OUT DMEM (Gibco, 10829-018), 15% KNOCK-OUT Serum Replacement (Gibco, 10828-028), GlutamaxX100, (Gibco, 35050-038), 1% Non-essential amino acids (NEAA) (Gibco, 11140-035), 0.1 mM b-mercaptoethanol (Sigma, M6250-250ML), penicillin–streptomycin (Biological Industries, 03-031-1B) and 4ul bFGF solution (10 μg/mL) per 10 mL (Peprotech, 100-18B), at 37 °C in a humidified incubator with 5% CO2. HT-29, MCF7 and HEK293T cell lines and were routinely cultured in DMEM containing 10% fetal bovine serum and 100 U penicillin/0.1 mg ml−1 streptomycin, at 37 °C in a humidified incubator with 5% CO2.

### Endodermal differentiation

Endodermal differentiation was performed as previously described (Loh et al. 2014). Pluripotent human stem cells were grown in the absence of MEF for four passages in mTeSR1 (StemCell Technologies, 05850) and seeded on Geltrex (invitrogen, A1413202). After 1-2 days of recovery in mTeSR1, hESC were washed with F12 (Gibco, 21765-029) to evacuate all mTeSR1 and then were treated for 24 hours with Activin A (100 ng/mL, R&D Systems, 338-AC-010), CHIR99021 (2 μM, Stemgent, 04-0004), and PI-103 (50 nM, Tocris, 2930) in CDM2 to specify APS. Afterwards, cells were washed (F12), then treated for 48 hours with Activin A (100 ng/mL) and LDN-193189/DM3189 (250 nM, Stemgent, 04-0074) in CDM2 to generate DE by day 3. Day 3 DE was patterned into AFG, PFG, or MHG by 4 days of continued differentiation in CDM2. DE was washed (F12), then differentiated as follows: AFG, A-83-01 (1 μM, Tocris, 2939) and LDN-193189 (250 nM, Stemgent, 04-0074); PFG, RA (2 μM, Sigma, R2625) and LDN-193189 (250 nM); MHG, BMP4 (10 ng/mL, R&D Systems, 314-BP-010), CHIR99021 (3 μM, Stemgent, 04-0004), and FGF2 (100 ng/mL, Peprotech, 100-18B), yielding day 7 anteroposterior domains. Media was refreshed every 24 hours for each differentiation step.

### HT-29 enterocytic differentiation

Enterocyte differentiation was performed as previously described (Zweibaum et al. 1985). HT-29 Cells were seeded in 90% confluence on ThInCerts (Greiner, 60-657641) in 6 well plates. Cells were cultured for 31 days in glucose free conditions (Sigma, 11966-025) and the medium was changed every 2 days.

### Transfections

Plasmid transfections for HEK293T, MCF7 and HT-29 were performed using PolyEthylene Imine (PEI) (PEI linear, M*r* 25,000, Polyscience). CRISPRi/a transient experiments were harvested after 72h. siRNAs were transfected into HT-29 cells at 10 nM siRNA pool or with control pool (Dharmacon) by using DharmaFect 4 (Dharmacon,T-2004-01/03) following the manufacturer’s protocol. Cells were harvested after 48h of siRNA treatment.

### Lentivirus production and stable lines generation

All lentivirus production was performed as previously described (Tiscornia et al. 2006). Medium was collected from plates 72 hr after transfection, filtered by VIVASPIN (Sartorius,VS2001), concentrated and stored –80°C. hESC and HT-29 cells were infected by lentiviral particles incubated in the growth medium containing and 8μg/ml Polybren (Sigma, 107689-10G) to attached cells, following selection after 24h for several passages for pool isolation.

### RNA and RT-qPCR

Total RNA was extracted from different cell lines and mouse tissues, by using RNeasy (Qiagen) according to the manufacturer’s protocol. cDNA was synthesized by using qScript Flex cDNA synthesis kit (Quanta, 95049). Fast SYBR Green master mix (Life, 4385614) was used for qPCR with gene-specific primers.

### Immunofluorescence

Cultured cells were fixed with 4% paraformaldehyde for 10 minutes. Fixated cells were permeabilized using 0.1% triton X-100 and blocked using 5% normal goat serum, incubated with a primary antibody, followed by incubation with a secondary antibody conjugated to a fluorescent dye. Antibodies used: Rabbit α-Eomes (Abcam, ab23345), Goat α-Sox17 (R&D Systems, AF1924), Goat α-Cdx2 (R&D Systems, AF3665), Goat α-Otx2 (R&D Systems, AF1979).

### Single-molecule FISH

Cultured cells were fixed with 4% paraformaldehyde 24 hr after plating. Tissue was frozen in Tissue-Tek O.C.T compound (Sakura 4583) blocks and sectioned using a Leica cryostat (CM3050) at 10 μm thickness. Libraries of 96 and 94 probes (**Table S2**) were designed to target human *HOXA-AS3* and mouse *Hoxaas3* RNA sequences, respectively and a commercially available library of 48 probes was used to detect HOXA5 (cat # VSMF-2538-5) (Stellaris RNA FISH probes, Biosearch Technologies). Hybridization conditions and imaging were as previously described (Itzkovitz et al. 2011; Lyubimova et al. 2013). smFISH imaging was performed on a Nikon-Ti-E inverted fluorescence microscope with a 100 × oil-immersion objective and a Photometrics Pixis 1024 CCD camera using MetaMorph software as previously described (Bahar Halpern and Itzkovitz 2016).

### CRISPR gRNA cloning

Guide RNAs were designed by CHOPCHOP. Cloning of plasmids was done following Zhang Lab General Protocol (http://www.genome-engineering.org/crispr/wp-content/uploads/2014/05/CRISPR-Reagent-Description-Rev20140509.pdf) by using pKLV-U6gRNA(BbsI)-PGKpuro2ABFP (Addgene plasmid #50946).

## Supplemental Tables

**Table S1: FANTOM5 expression data**

**Table S2: smFISH probes**

**Table S3: Markers used for validation**

**Table S4: Primers and siRNAs**

## Author contributions

ND and IU conceived the study. ND carried out the experiments. ND and IU analyzed data. ND and IU wrote the manuscript. EA assisted with design and analysis of the hESC experiments.

## References

Bahar Halpern K, Itzkovitz S. 2016. Single molecule approaches for quantifying transcription and degradation rates in intact mammalian tissues. Methods 98: 134–142.

Cabili MN, Trapnell C, Goff L, Koziol M, Tazon-Vega B, Regev A, Rinn JL. 2011. Integrative annotation of human large intergenic noncoding RNAs reveals global properties and specific subclasses. Genes Dev 25: 1915–1927.

Casaca A, Hauswirth GM, Bildsoe H, Mallo M, McGlinn E. 2018. Regulatory landscape of the Hox transcriptome. Int J Dev Biol 62: 693–704.

Clevers H. 2013. The intestinal crypt, a prototype stem cell compartment. Cell 154: 274–284.

Consortium, Fantom, The RP, Clst, Forrest AR, Kawaji H, Rehli M, Baillie JK, de Hoon MJ, Lassmann T, Itoh M, et al. 2014. A promoter-level mammalian expression atlas. Nature 507: 462–470.

Deschamps J, van Nes J. 2005. Developmental regulation of the Hox genes during axial morphogenesis in the mouse. Development 132: 2931–2942.

Gilbert LA, Larson MH, Morsut L, Liu Z, Brar GA, Torres SE, Stern-Ginossar N, Brandman O, Whitehead EH, Doudna JA, et al. 2013. CRISPR-mediated modular RNA-guided regulation of transcription in eukaryotes. Cell 154: 442–451.

Gil N, Ulitsky I. 2020. Regulation of gene expression by cis-acting long non-coding RNAs. Nat Rev Genet 21: 102–117.

Grainger S, Savory JGA, Lohnes D. 2010. Cdx2 regulates patterning of the intestinal epithelium. Dev Biol 339: 155–165.

Greer JM, Puetz J, Thomas KR, Capecchi MR. 2000. Maintenance of functional equivalence during paralogous Hox gene evolution. Nature 403: 661–665.

Guttman M, Amit I, Garber M, French C, Lin MF, Feldser D, Huarte M, Zuk O, Carey BW, Cassady JP, et al. 2009. Chromatin signature reveals over a thousand highly conserved large non-coding RNAs in mammals. Nature 458: 223–227.

Haber AL, Biton M, Rogel N, Herbst RH, Shekhar K, Smillie C, Burgin G, Delorey TM, Howitt MR, Katz Y, et al. 2017. A single-cell survey of the small intestinal epithelium. Nature 551: 333–339.

Hezroni H, Koppstein D, Schwartz MG, Avrutin A, Bartel DP, Ulitsky I. 2015. Principles of Long Noncoding RNA Evolution Derived from Direct Comparison of Transcriptomes in 17 Species. Cell Rep. http://dx.doi.org/10.1016/j.celrep.2015.04.023.

Hoegg S, Meyer A. 2005. Hox clusters as models for vertebrate genome evolution. Trends Genet 21: 421–424.

Hornstein E, Mansfield JH, Yekta S, Hu JK-H, Harfe BD, McManus MT, Baskerville S, Bartel DP, Tabin CJ. 2005. The microRNA miR-196 acts upstream of Hoxb8 and Shh in limb development. Nature 438: 671–674.

Huang H-H, Chen F-Y, Chou W-C, Hou H-A, Ko B-S, Lin C-T, Tang J-L, Li C-C, Yao M, Tsay W, et al. 2019. Long non-coding RNA HOXB-AS3 promotes myeloid cell proliferation and its higher expression is an adverse prognostic marker in patients with acute myeloid leukemia and myelodysplastic syndrome. BMC Cancer 19: 617.

Huang J-Z, Chen M, Chen D, Gao X-C, Zhu S, Huang H, Hu M, Zhu H, Yan G-R. 2017. A Peptide Encoded by a Putative lncRNA HOXB-AS3 Suppresses Colon Cancer Growth. Mol Cell 68: 171–184.e6.

Ihry RJ, Worringer KA, Salick MR, Frias E, Ho D, Theriault K, Kommineni S, Chen J, Sondey M, Ye C, et al. 2018. p53 inhibits CRISPR–Cas9 engineering in human pluripotent stem cells. Nat Med 24: 939–946.

Itzkovitz S, Lyubimova A, Blat IC, Maynard M, van Es J, Lees J, Jacks T, Clevers H, van Oudenaarden A. 2011. Single-molecule transcript counting of stem-cell markers in the mouse intestine. Nat Cell Biol 14: 106–114.

Izpisúa-Belmonte JC, Tickle C, Dollé P, Wolpert L, Duboule D. 1991. Expression of the homeobox Hox-4 genes and the specification of position in chick wing development. Nature 350: 585–589.

Kanai-Azuma M, Kanai Y, Gad JM, Tajima Y, Taya C, Kurohmaru M, Sanai Y, Yonekawa H, Yazaki K, Tam PPL, et al. 2002. Depletion of definitive gut endoderm in Sox17-null mutant mice. Development 129: 2367–2379.

Kashyap V, Gudas LJ, Brenet F, Funk P, Viale A, Scandura JM. 2011. Epigenomic reorganization of the clustered Hox genes in embryonic stem cells induced by retinoic acid. J Biol Chem 286: 3250–3260.

Kmita M, Tarchini B, Zàkàny J, Logan M, Tabin CJ, Duboule D. 2005. Early developmental arrest of mammalian limbs lacking HoxA/HoxD gene function. Nature 435: 1113–1116.

Kumar N, Tsai Y-H, Chen L, Zhou A, Banerjee KK, Saxena M, Huang S, Toke NH, Xing J, Shivdasani RA, et al. 2019. The lineage-specific transcription factor CDX2 navigates dynamic chromatin to control distinct stages of intestine development. Development 146. http://dx.doi.org/10.1242/dev.172189.

Lin MF, Jungreis I, Kellis M. 2011. PhyloCSF: a comparative genomics method to distinguish protein coding and non-coding regions. Bioinformatics 27: i275–82.

Lin S, Zhang R, An X, Li Z, Fang C, Pan B, Chen W, Xu G, Han W. 2019. LncRNA HOXA-AS3 confers cisplatin resistance by interacting with HOXA3 in non-small-cell lung carcinoma cells. Oncogenesis 8: 60.

Loh KM, Ang LT, Zhang J, Kumar V, Ang J, Auyeong JQ, Lee KL, Choo SH, Lim CYY, Nichane M, et al. 2014. Efficient endoderm induction from human pluripotent stem cells by logically directing signals controlling lineage bifurcations. Cell Stem Cell 14: 237–252.

Lyubimova A, Itzkovitz S, Junker JP, Fan ZP, Wu X, van Oudenaarden A. 2013. Single-molecule mRNA detection and counting in mammalian tissue. Nat Protoc 8: 1743–1758.

Mansfield JH, Harfe BD, Nissen R, Obenauer J, Srineel J, Chaudhuri A, Farzan-Kashani R, Zuker M, Pasquinelli AE, Ruvkun G, et al. 2004. MicroRNA-responsive’sensor’transgenes uncover Hox-like and other developmentally regulated patterns of vertebrate microRNA expression. Nat Genet 36: 1079–1083.

Martínez-Maqueda D, Miralles B, Recio I. 2015. HT29 Cell Line. The Impact of Food Bioactives on Health 113–124. http://dx.doi.org/10.1007/978-3-319-16104-4_11.

Mercer TR, Dinger ME, Sunkin SM, Mehler MF, Mattick JS. 2008. Specific expression of long noncoding RNAs in the mouse brain. Proc Natl Acad Sci U S A 105: 716–721.

Papaioannou D, Petri A, Dovey OM, Terreri S, Wang E, Collins FA, Woodward LA, Walker AE, Nicolet D, Pepe F, et al. 2019. The long non-coding RNA HOXB-AS3 regulates ribosomal RNA transcription in NPM1-mutated acute myeloid leukemia. Nat Commun 10: 5351.

Perry RB-T, Ulitsky I. 2016. The functions of long noncoding RNAs in development and stem cells. Development 143: 3882–3894.

Pijuan-Sala B, Griffiths JA, Guibentif C, Hiscock TW, Jawaid W, Calero-Nieto FJ, Mulas C, Ibarra-Soria X, Tyser RCV, Ho DLL, et al. 2019. A single-cell molecular map of mouse gastrulation and early organogenesis. Nature 566: 490–495.

Qi LS, Larson MH, Gilbert LA, Doudna JA, Weissman JS, Arkin AP, Lim WA. 2013. Repurposing CRISPR as an RNA-guided platform for sequence-specific control of gene expression. Cell 152: 1173–1183.

Rinn JL, Kertesz M, Wang JK, Squazzo SL, Xu X, Brugmann SA, Goodnough LH, Helms JA, Farnham PJ, Segal E, et al. 2007. Functional demarcation of active and silent chromatin domains in human HOX loci by noncoding RNAs. Cell 129: 1311–1323.

Rousset M. 1986. The human colon carcinoma cell lines HT-29 and Caco-2: Two in vitro models for the study of intestinal differentiation. Biochimie 68: 1035–1040. http://dx.doi.org/10.1016/s0300-9084(86)80177-8.

Sarropoulos I, Marin R, Cardoso-Moreira M, Kaessmann H. 2019. Developmental dynamics of lncRNAs across mammalian organs and species. Nature 571: 510–514.

Smillie CS, Biton M, Ordovas-Montanes J, Sullivan KM, Burgin G, Graham DB, Herbst RH, Rogel N, Slyper M, Waldman J, et al. 2019. Intra- and Inter-cellular Rewiring of the Human Colon during Ulcerative Colitis. Cell 178: 714–730.e22.

Soshnikova N, Duboule D. 2008. Epigenetic regulation of Hox gene activation: the waltz of methyls. Bioessays 30: 199–202.

Soshnikova N, Duboule D. 2009. Epigenetic regulation of vertebrate Hox genes: a dynamic equilibrium. Epigenetics 4: 537–540.

Tabula Muris Consortium, Overall coordination, Logistical coordination, Organ collection and processing, Library preparation and sequencing, Computational data analysis, Cell type annotation, Writing group, Supplemental text writing group, Principal investigators. 2018. Single-cell transcriptomics of 20 mouse organs creates a Tabula Muris. Nature 562: 367–372.

Tiscornia G, Singer O, Verma IM. 2006. Production and purification of lentiviral vectors. Nature Protocols 1: 241–245. http://dx.doi.org/10.1038/nprot.2006.37.

Tyler DM, Okamura K, Chung W-J, Hagen JW, Berezikov E, Hannon GJ, Lai EC. 2008. Functionally distinct regulatory RNAs generated by bidirectional transcription and processing of microRNA loci. Genes Dev 22: 26–36.

Ulitsky I. 2016. Evolution to the rescue: using comparative genomics to understand long non-coding RNAs. Nat Rev Genet. http://dx.doi.org/10.1038/nrg.2016.85.

Varlakhanova N, Cotterman R, Bradnam K, Korf I, Knoepfler PS. 2011. Myc and Miz-1 have coordinate genomic functions including targeting Hox genes in human embryonic stem cells. Epigenetics Chromatin 4: 20.

Verzi MP, Hatzis P, Sulahian R, Philips J, Schuijers J, Shin H, Freed E, Lynch JP, Dang DT, Brown M, et al. 2010. TCF4 and CDX2, major transcription factors for intestinal function, converge on the same cis-regulatory regions. Proc Natl Acad Sci U S A 107: 15157–15162.

Wang KC, Yang YW, Liu B, Sanyal A, Corces-Zimmerman R, Chen Y, Lajoie BR, Protacio A, Flynn RA, Gupta RA, et al. 2011. A long noncoding RNA maintains active chromatin to coordinate homeotic gene expression. Nature 472: 120–124.

Wong SFL, Agarwal V, Mansfield JH, Denans N, Schwartz MG, Prosser HM, Pourquié O, Bartel DP, Tabin CJ, McGlinn E. 2015. Independent regulation of vertebral number and vertebral identity by microRNA-196 paralogs. Proc Natl Acad Sci U S A 112: E4884–93.

Wu F, Zhang C, Cai J, Yang F, Liang T, Yan X, Wang H, Wang W, Chen J, Jiang T. 2017. Upregulation of long noncoding RNA HOXA-AS3 promotes tumor progression and predicts poor prognosis in glioma. Oncotarget 8: 53110–53123.

Yekta S, Tabin CJ, Bartel DP. 2008. MicroRNAs in the Hox network: an apparent link to posterior prevalence. Nat Rev Genet 9: 789–796.

Zhang H, Liu Y, Yan L, Zhang M, Yu X, Du W, Wang S, Li Q, Chen H, Zhang Y, et al. 2018. Increased levels of the long noncoding RNA, HOXA-AS3, promote proliferation of A549 cells. Cell Death Dis 9: 707.

Zhu X, Chen D, Liu Y, Yu J, Qiao L, Lin S, Chen D, Zhong G, Lu X, Wang Y, et al. 2019. Long Noncoding RNA HOXA-AS3 Integrates NF-κB Signaling To Regulate Endothelium Inflammation. Mol Cell Biol 39. http://dx.doi.org/10.1128/MCB.00139-19.

Zhu X-X, Yan Y-W, Chen D, Ai C-Z, Lu X, Xu S-S, Jiang S, Zhong G-S, Chen D-B, Jiang Y-Z. 2016. Long non-coding RNA HoxA-AS3 interacts with EZH2 to regulate lineage commitment of mesenchymal stem cells. Oncotarget 7: 63561–63570.

Zweibaum A, Pinto M, Chevalier G, Dussaulx E, Triadou N, Lacroix B, Haffen K, Brun J-L, Rousset M. 1985. Enterocytic differentiation of a subpopulation of the human colon tumor cell line HT-29 selected for growth in sugar-free medium and its inhibition by glucose. Journal of Cellular Physiology 122: 21–29. http://dx.doi.org/10.1002/jcp.1041220105.

